# “Sex-divergent responses to microglial depletion suggest distinct regulatory dependencies in the aged brain”

**DOI:** 10.64898/2026.09.03.745468

**Authors:** Amila Beganovic, Matthias Flotho, Jinxiu Li, Ian H. Guldner, Marvin Reich, Vanessa Wahl, Simon Graf, Philipp Donate, Shusruto Rishik, Esugen D. Dashdorj, Nicole Ludwig, Rolf Müller, Tal Iram, Tony Wyss-Coray, Viktoria Wagner, Andreas Keller

## Abstract

Microglia are essential for brain homeostasis, yet their roles in the aged brain remain poorly defined. Using microRNA (miRNA) profiling, cellular-resolution spatial transcriptomics, and bulk proteomics in 21-month-old mice, we characterize sex-dimorphic responses to microglial depletion via CSF1R inhibition (PLX5622 treatment). Microglia-enriched miRNAs, notably miR-146a-5p and miR-223-3p, were downregulated across different brain regions in both sexes. Transcriptional responses were sex dimorphic: females showed predominantly cell-type-specific downregulation, while males showed bidirectional changes including upregulation of *Lzts3*, *Shank3*, and *Fgfbp1* alongside downregulation of *Ang*. Proteomic changes were larger in magnitude and independent from mRNA changes: males exhibited 295 differentially expressed proteins (DEPs) versus 34 in females (8.7-fold difference). Male DEPs had opposing directional shifts in synaptic vesicle proteins (upregulated) and mitochondrial ATP synthesis machinery (downregulated). These data describe sex-dimorphic molecular consequences of microglial loss in the aged brain and identify candidate post-transcriptional mechanisms warranting further investigation.

## Introduction

Brain aging represents a fundamental biological process characterized by progressive molecular and cellular changes that compromise neural function and increase susceptibility to neurodegenerative diseases^1^. This aging process is characterized by striking regional selectivity in vulnerability to age-associated changes^2^. Moreover, emerging evidence suggests that brain aging processes are sexually dimorphic^3^, with males and females exhibiting distinct molecular responses to age-related stressors, yet the mechanistic basis for these sex differences remains poorly understood.

MicroRNAs (miRNAs) have emerged as key post-transcriptional regulators exerting control of mRNA stability, translation and downstream protein abundance^4^. Comprehensive spatio-temporal atlases have identified sex-independent, age-related changes in microglial-enriched miRNAs across multiple brain regions, including miR-146a-5p, among a small set of consistently altered brain aging miRNAs^5,6^. However, whether microglia-enriched miRNA programs functionally contribute to sex-biased molecular outcomes during brain aging, particularly at the transcriptomic and proteomic levels, remains unexplored.

Colony-stimulating factor 1 receptor (CSF1R) is a class III receptor tyrosine kinase activated by two ligands, CSF1 and IL-34^7^. Ligand binding triggers downstream PI3K-AKT signaling that sustains microglial survival, MAPK/ERK signaling that drives proliferation, and Src-family kinase activity that supports differentiation and maintenance^8^. Adult microglia depend on continuous CSF1R signaling for viability, with regional sensitivity reflecting differential CSF1 and IL-34 expression in white versus gray matter^7,9^. In the aged brain, microglia remain dependent on CSF1R signaling for survival, and CSF1R inhibition reduces age-associated neuroinflammation and cellular senescence while improving cognitive performance in aged mice^10,11^. CSF1R signaling is conserved between mice and humans, with CSF1R inhibitors now in clinical development for neurodegenerative diseases^12^. The aged mouse allows microglial number to be selectively and reversibly manipulated in vivo, isolating microglia-dependent contributions to the sex-biased vulnerability seen in human neurodegeneration.

Pharmacological inhibition of CSF1R using small-molecule inhibitors such as PLX3397 and PLX5622 has emerged as the primary approach for selective microglia depletion in the adult brain^9,13^. In young healthy mice, microglia appear largely dispensable for basic neural function^9^. However, in disease and aging contexts, microglia perturbation has revealed critical roles in Alzheimer’s pathology^14,15^, traumatic brain injury^16^, and radiation-induced cognitive decline^17^. Recent work further demonstrates that partial microglia depletion, reducing numbers to young-like levels, may improve synaptic plasticity while avoiding risks of complete ablation^11^, and that depletion produces dose-dependent reductions in neuroinflammation and cellular senescence markers in aged brains^10^. However, the molecular consequences of microglial depletion across the broader cellular system of the aged brain remain incompletely characterized, particularly with respect to potential sex-dependent effects.

Microglia exhibit intrinsic sex differences in transcriptomic and proteomic profiles that persist even after transplantation into opposite-sex brains^18^. Critically, the microglia response to CSF1R inhibition itself is sex-dimorphic: depletion efficiency, survival signaling pathways, and functional consequences differ between males and females^19,20^. Together, these findings demonstrate that microglial depletion elicits sex-dimorphic responses at the cellular level. Whether these differences extend to downstream molecular consequences, spanning miRNA mediated regulation, transcriptional remodeling, and proteomics adaptation, has not been characterized.

Given microglia’s role in age-related neuroinflammation and miRNA dysregulation, understanding how the aged brain depends on ongoing microglial activity represents an important open question. Microglial depletion provides a trackable perturbation to assess the functional consequences of microglial loss in an already-aged system – revealing which molecular programs require continuous microglial support to be maintained, and whether this dependency differs between sexes.

The specific objective of this study was to determine whether microglial depletion in the aged brain produces sex-dimorphic molecular consequences, and if so, to identify at which regulatory layer(s) these differences emerge. We hypothesized that microglial depletion would produce sex-differential molecular signatures, without predicting a priori which layer or direction. To address this, we integrated bulk miRNA sequencing, spatial transcriptomics and bulk proteomics as a comprehensive approach to dissecting sex-specific brain aging mechanisms following CSF1R inhibition in 21-month-old mice. While bulk miRNA and proteomic analyses capture system-wide regulatory changes and their downstream functional consequences, spatial transcriptomics reveals the cellular and regional context of these alterations. This integrated framework maps candidate miRNA-mRNA-protein networks and their spatial distribution across brain regions. By applying this integrated framework to aged mice, we identify sex-dimorphic consequences of microglial loss that extend current understanding of microglial roles in brain aging and highlight the potential relevance of biological sex in microglial-targeting therapeutic contexts.

## Results

### Microglial depletion in aged mice preserves non-microglial cell populations across brain regions in both sexes

Microglia regulate neuronal health in aging and neurodegeneration, yet sex-specific microglial functions in the aged brain remain poorly understood^21^. To investigate sex-specific responses to microglial loss, we depleted microglia in 21-month-old C57BL/6J mice using the colony-stimulating factor 1 receptor (CSF1R) inhibitor PLX5622 for three weeks (Control: n=4 per sex; PLX: n=6 per sex). We performed integrated multi-omics profiling across the same cohort, including small RNA sequencing from four anatomically and functionally different brain regions (olfactory bulb, caudate putamen, bregma -2.06, and cerebellum), cellular-resolution spatial transcriptomics on the bregma -2.06 coronal section using Stereo-seq (∼2.5 million cells across 20 sections), and bulk proteomics on sections posterior to bregma -2.06 (**Fig. 1A**). Each omic layer was analyzed independently to establish sex-specific responses before integration to reveal convergent biological mechanisms.

**Figure 1:**
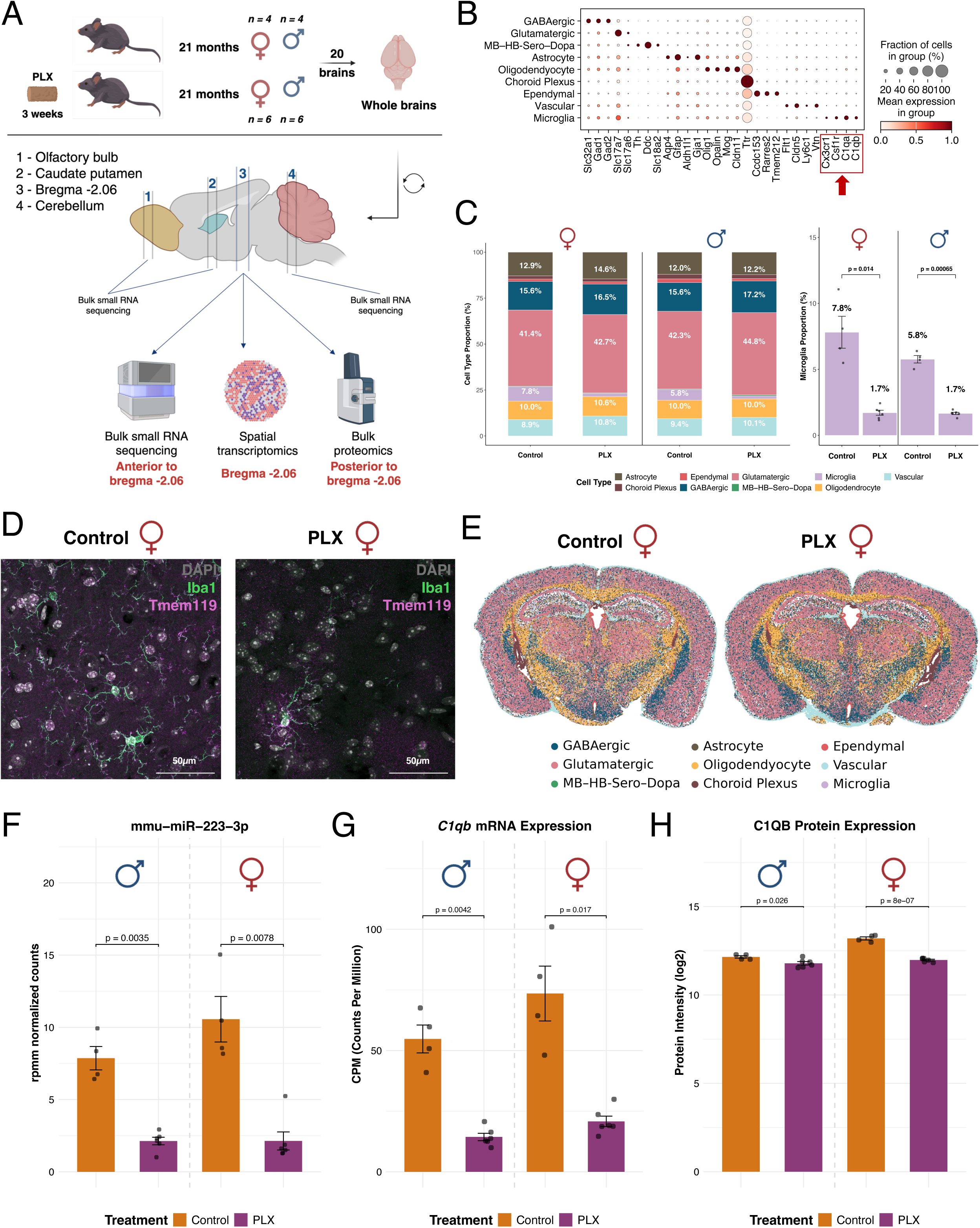
Microglial depletion in aged mice validated across miRNA, transcriptome, and proteome levels. A. Experimental design. 21-month-old C57BL/6J mice (n = 4/sex control; n = 6/sex PLX) received control or PLX5622-containing chow (1200ppm) for 3 weeks. Brain tissue was analyzed by bulk small RNA sequencing across four brain regions (1: olfactory bulb; 2: caudate putamen; 3: bregma -2.06; 4: cerebellum), spatial transcriptomics (bregma -2.06 coronal section), and bulk proteomics (10µm coronal sections immediately posterior to the bregma -2.06 section). Labels beneath the three platforms indicate relative position of the serial 10µm coronal sections used at the bregma -2.06 level. B. Dot plot showing mean expression of cell type marker genes across six major populations identified by spatial transcriptomics. Dot size indicates fraction of cells expressing each marker; color indicates mean expression level within the group. Color indicates per-gene min-max normalized mean expression, which linearly rescales each gene’s group mean across cell populations to the interval [0, 1]. Red box highlights microglial markers (*Cx3cr1*, *Csf1r*, *C1qa*, *C1qb*). C. Stacked bar plot showing cell type proportions across experimental groups (left). Cell type composition remained stable following PLX treatment in both sexes, with no compensatory proliferation of non-microglial populations. Inset bar plots (right) highlight microglial proportions specifically. D. Representative confocal microscopy images (40x magnification) of the motor cortex region from 21-month-old female control (left) and PLX-treated (right) mice. Brain sections were stained for Iba1 (green) and Tmem119 (magenta), microglia-specific markers, and counterstained with DAPI (gray) to visualize nuclei. E. Spatial mapping of cell type distribution of all cell types in control and PLX-treated females. F. MiR-223-3p expression (rpmm normalized counts) in the bregma -2.06 coronal section measured by small RNA sequencing. Statistical significance determined by Welch’s t-test comparing Control vs. PLX within each sex. G. *C1qb* mRNA expression (counts per million, CPM) from spatial transcriptomics. Statistical significance determined by Welch’s t-test comparing Control vs. PLX within each sex. H. C1QB protein intensity (log2) from bulk proteomics. Statistical significance determined by Student’s t-test (Perseus) comparing Control vs. PLX within each sex.

We annotated nine cell populations in the spatial transcriptomics data based on established marker genes^22^ (**Fig. 1B**). Six major populations, namely astrocytes (*Aqp4, Gfap, Aldh1l1, Gja1*), GABAergic neurons (*Gad1/2, Slc32a1*), glutamatergic neurons (*Slc17a7, Slc17a6*), oligodendrocytes (*Olig1, Cldn11*), vascular cells (*Cldn5, Flt1*), and microglia (*Cx3cr1, C1qa, C1qb, Csf1r*), comprised a mean of 96.6% of total cells and were the focus of downstream analyses. PLX treatment depleted microglia by 79% in females and 71% in males (**Fig. 1C**). This depletion efficiency is consistent with previous reports in aged mice, where PLX typically achieves 69-84% microglial reduction^10^. We confirmed depletion by immunohistochemical staining demonstrating reduced Iba1+ and Tmem119+ cell density across multiple brain regions including motor cortex, cerebellum, and thalamus (**Fig. 1D, Supp. Fig. 1A-C**), together with reduced CD206+ signal marking border-associated macrophages (**Supp. Fig. 2A-C**). CD31+ vascular staining showed no substantial reduction (**Supp. Fig. 2A-C**), indicating the reduction in CD206+ cells reflects macrophage loss rather than vascular disruption. Non-microglial cell-type proportions remained stable (**Fig. 1C, Supp. Table 1**), consistent with CSF1R enrichment in microglia within the central nervous system (CNS)^9^. The only exception was a modest reduction in ependymal cells in males (mean reduction of 0.76%), likely reflecting co-capture of CSF1R-dependent choroid plexus macrophages, which reside on the surface of the ependymal compartment lining the ventricles^23^. Spatial mapping confirmed preserved anatomical organization of major cell populations across all samples following microglial depletion (**Fig. 1E, Supp. Fig. 3A-D**).

**Figure 2:**
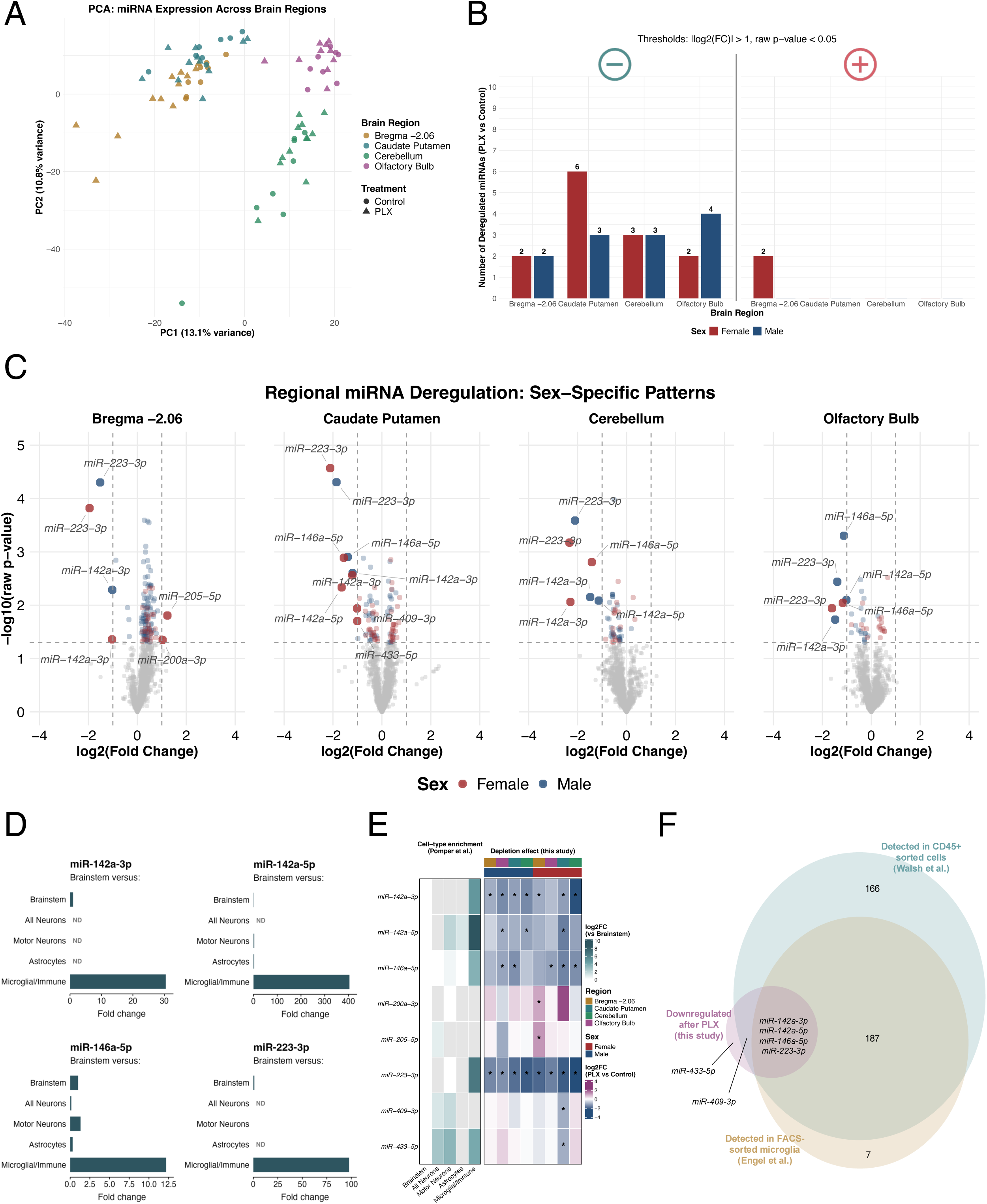
Regional miRNA dysregulation following microglial depletion reveals conserved microglial-enriched signatures. A. Principal component analysis (PCA) of miRNA expression across four brain regions (olfactory bulb, caudate putamen, bregma -2.06 coronal section, cerebellum). Shape indicates treatment; colors indicate brain region. PC1 (13.1% variance) primarily separated by anatomical regions. B. Bar plot showing the number of deregulated miRNAs per brain region and sex. Differential expression was determined by |log2FC| > 1.0, and raw p-value < 0.05 (FDR adjusted results in **Supp. Fig. 4E**). Numbers indicate count of downregulated and upregulated miRNAs (colors indicate sex). C. Volcano plots of miRNA differential expression in four brain regions, separated by sex. Labeled miRNAs represent microglial-enriched miRNAs (miR-142a-3p, miR-142a-5p, miR-146a-5p and miR-223-3p) and non-microglial miRNAs (miR-200-3p, miR-205-5p, miR-409-3p and miR-433-5p), classified by relative expression across sorted CNS cell populations reported in published miRNA profiling database^24,30^. Differential expression was determined by |log2FC| > 1.0, and raw p-value < 0.05. D. Bar plots showing cell-type enrichment of the four convergently downregulated miRNAs (miR-142a-3p, miR-142a-5p, miR-146a-5p, and miR-223-3p), based on CNS cell-type expression profiles from^24,30^. Bars indicate fold change of each miRNA in a given brain cell type relative to brainstem. All four miRNAs show strong microglial/immune enrichment. E. Heatmap of six miRNAs deregulated following CSF1R inhibition. Left: log2FC in expression per CNS cell type relative to brainstem from Pomper et al. database^24,30^. Right: log2FC for PLX versus control within each region and sex. Asterisks denote |log2FC| > 1.0, and raw p-value < 0.05. Both blocks have a log2FC color scale, for visual comparison only (the two FC types arise from different types of comparisons). F. Euler diagram showing overlap of depletion-associated miRNAs identified in this study with microglia-associated miRNAs from two independent datasets: miRNAs expressed in FACS-sorted microglia^5^ and miRNAs expressed from bulk CD45+ sorted cells^6^. The overlap highlights the conserved microglial identity of the miRNAs lost following depletion.

To further validate microglial depletion across platforms, we examined representative microglial markers in each omic layer. MiR-223-3p, a microglial-enriched miRNA that increases in aged microglia^5,24^, decreased in both sexes following PLX treatment (**Fig. 1F**). For transcriptome and proteome validation, we chose complement component 1q, which shows near-exclusive microglial expression in the CNS and serves as one of the most specific and abundant microglial products^25^. Spatial transcriptomics demonstrates a marked reduction in complement component *C1qb* in both sexes (72% in females and 74% in males) (**Fig. 1G**). Bulk proteomics showed more modest C1QB reduction (approximately 10%; p < 0.05) (**Fig. 1H**). This transcript-protein discordance likely reflects two factors: the extended half-life of extracellular complement proteins and continued production by residual microglia. Furthermore, it illustrates that protein abundance does not always directly track mRNA levels, particularly for secreted or long-lived proteins^26^, a principle critical for interpreting subsequent proteomic findings. Multi-omics validation confirmed effective microglial depletion across miRNA, transcriptome and proteome levels while maintaining stable non-microglial cell type proportions.

### Microglial depletion drives region-specific miRNA dysregulation dominated by loss of microglia-enriched miRNAs

Post-transcriptional gene regulation through miRNAs represents a critical but understudied mechanism by which microglia may influence brain homeostasis. To identify altered miRNAs following microglial depletion, we performed bulk small RNA sequencing across four different brain regions. Quality control confirmed consistent library composition with miRNAs representing an average of 52.4% of mapped reads (followed by lncRNA at 29%) and high inter-sample correlations (Spearman > 0.85), indicating strong technical reproducibility (**Supp. Fig. 4A-B**). Principal component analysis (PCA) of miRNA expression revealed that anatomical region explains more variance than treatment or sex (**Fig. 2A, Supp. Fig. 4A, C-D**). This regional identity dominance is consistent with a spatio-temporal atlas of mouse brain miRNA expression, in which anatomical region accounted for approximately 54% of total miRNA variance compared to <1% for age as an independent factor^5^. Because regional baseline differences exceed treatment- and sex-induced changes in absolute magnitude, pooling samples across regions would dilute the depletion signal. We therefore stratified all differential expression analyses by region and sex, assessing treatment effects within each region against region-matched controls.

Given the exploratory nature of this study, we used raw p-values for primary differential expression analysis, with adjusted p-value results reported alongside in supplementary materials. With microglial depletion, a total of 25 miRNAs were downregulated across regions and sexes and two miRNAs were upregulated in female bregma -2.06 (raw p < 0.05; |log2FC| > 1.0) (**Fig. 2B, Supp. Fig. 4E**; complete list of deregulated miRNAs in **Supp. Table 2**). The caudate putamen showed the greatest number of deregulated miRNAs (6 downregulated miRNAs in females; 3 in males), followed by olfactory bulb (2 in females; 4 in males), cerebellum (3 per sex), and bregma -2.06 (2 per sex). This regional hierarchy may reflect differences in microglial abundance or depletion efficiency across brain regions^27^. The bregma -2.06 coronal level was selected for integrated multi-omics analysis as it simultaneously captures multiple anatomically and functionally distinct regions (cortex, hippocampus, striatum and thalamus), within a single tissue plane, maximizing biological interpretability of regional comparisons^28^.

The only upregulated miRNAs were miR-200a-3p and miR-205-5p (raw p < 0.05; log2FC > 1.0) in females at the bregma -2.06 level (**Fig. 2B-C**). Cell-type-resolved profiling confirmed neither miRNA is microglia-enriched (**Supp. Fig. 4F**), indicating their upregulation is likely not caused by microglia depletion. Whether this female-specific upregulation represents a secondary consequence of microglial loss, or a depletion-independent sex effect remains to be determined. Building on the shared downregulation patterns identified above, we examine region-specific depletion signatures in detail (**Fig. 2C**). Notably, bregma -2.06 section shared substantially overlapping miRNA signatures with other brain regions. MiR-223-3p was downregulated in all four brain regions in both sexes, and it was the only miRNA to retain significance after multiple testing correction (in males at bregma -2.06 and caudate putamen, and in females at caudate putamen) (adjusted p < 0.05; |log2FC| > 1.0). This identifies miR-223-3p as the miRNA showing the broadest regional consistency of downregulation across the dataset. MiR-142a-3p was downregulated in all regions except female olfactory bulb, while miR-146a-5p was not downregulated in bregma -2.06 (both sexes) or male cerebellum. Two additional miRNAs, miR-409-3p and miR-433-5p, were downregulated uniquely only in females in the caudate putamen, the brain region dependent on neuron-to-microglia IL-34/CSF1R signaling for microglial maintenance^7,29^.

The convergent downregulation of miR-142a-3p/5p, miR-146a-5p, miR-223-3p across multiple brain regions raised the question of whether these signatures share a common cellular origin. We used a publicly available database of cell-type-enriched miRNA expression in the mouse CNS, built from Cre-dependent Argonaute-2 (Ago2) affinity-purification data, and confirmed that all four miRNAs are strongly enriched in the microglia/immune compartment relative to brainstem, neurons, motor neurons and astrocytes (**Fig. 2D, Supp. Table 3**)^24,30^. Of the eight miRNAs deregulated across any region or sex, only these four were enriched in microglia/immune populations in independent reference datasets (**Fig. 2E, Supp. Fig. 4F-G**), supporting their microglial origin.

To assess whether our PLX-downregulated miRNAs are detected in microglia-enriched populations, we performed a three-way comparison against two external microglia miRNA reference datasets (**Fig. 2F**). These include miRNAs detected in aged versus young FACS-sorted microglia from Engel et al. (84 weeks (W) vs 12 weeks (W))^5^ and in the CD45+ immune fraction relative to bulk CNS tissue from Walsh et al.^6^. Four miRNAs - miR-142a-3p, miR-142a-5p, miR-146a-5p, and miR-223-3p – were detected across all three datasets, providing convergent evidence from independent platforms that these miRNAs are expressed in microglia and are sensitive to microglial perturbation. MiR-409-3p shows neuronal expression in cell-type-resolved CNS profiles (**Supp. Fig. 4G**)^24,30^ and was shared between our dataset and the Walsh et al. CD45+ fraction. We interpret its downregulation as a possible secondary neuronal response to disrupted microglia-neuron crosstalk rather than direct depletion of microglia-enriched transcripts. MiR-433-5p was unique to our depletion set. However, existing cell-type-resolved reference data do not clearly resolve its cellular origin. We therefore do not attribute its downregulation to a specific cellular source. In contrast, miR-155-5p, another canonical microglial inflammatory miRNA^31^, was not deregulated in any region in our datasets, but it was detected as deregulated in the other two datasets^5,6^ (**Fig. 2F**). This suggests that not all microglial miRNAs are sensitive to acute CSF1R inhibition, likely consistent with low baseline expression in aged homeostatic microglia or preferential expression in the PLX-resistant microglial subpopulations that persist after treatment.

Together, these findings establish that microglial depletion in the aged brain preferentially eliminates microglia-enriched miRNAs supported by cross-dataset evidence (**Fig. 2D-F**), alongside a smaller number of non-microglial miRNAs whose downregulation likely reflects secondary responses to microglial loss. Having characterized these miRNA-level changes, we next examined cellular-resolution transcriptomic responses to determine how individual cell types and brain regions respond to microglial loss.

### Sex-dimorphic transcriptional responses to microglial depletion reveal differential cellular vulnerability

Sex-stratified differential expression analysis (PLX vs Control) across six major cell types revealed differing transcriptional susceptibility (**Fig. 3A**). We excluded genes with predominant microglial expression signatures and genes lacking reliable cell-type annotation (DEGs counts prior to this filtering are provided in **Supp. Fig. 5A-C**). This filtering step corrected for a technical limitation of Stereo-seq spatial transcriptomics, wherein microglia transcripts co-captured neighboring bins can be misattributed as treatment-induced transcriptional responses in non-microglial cell types.

**Figure 3:**
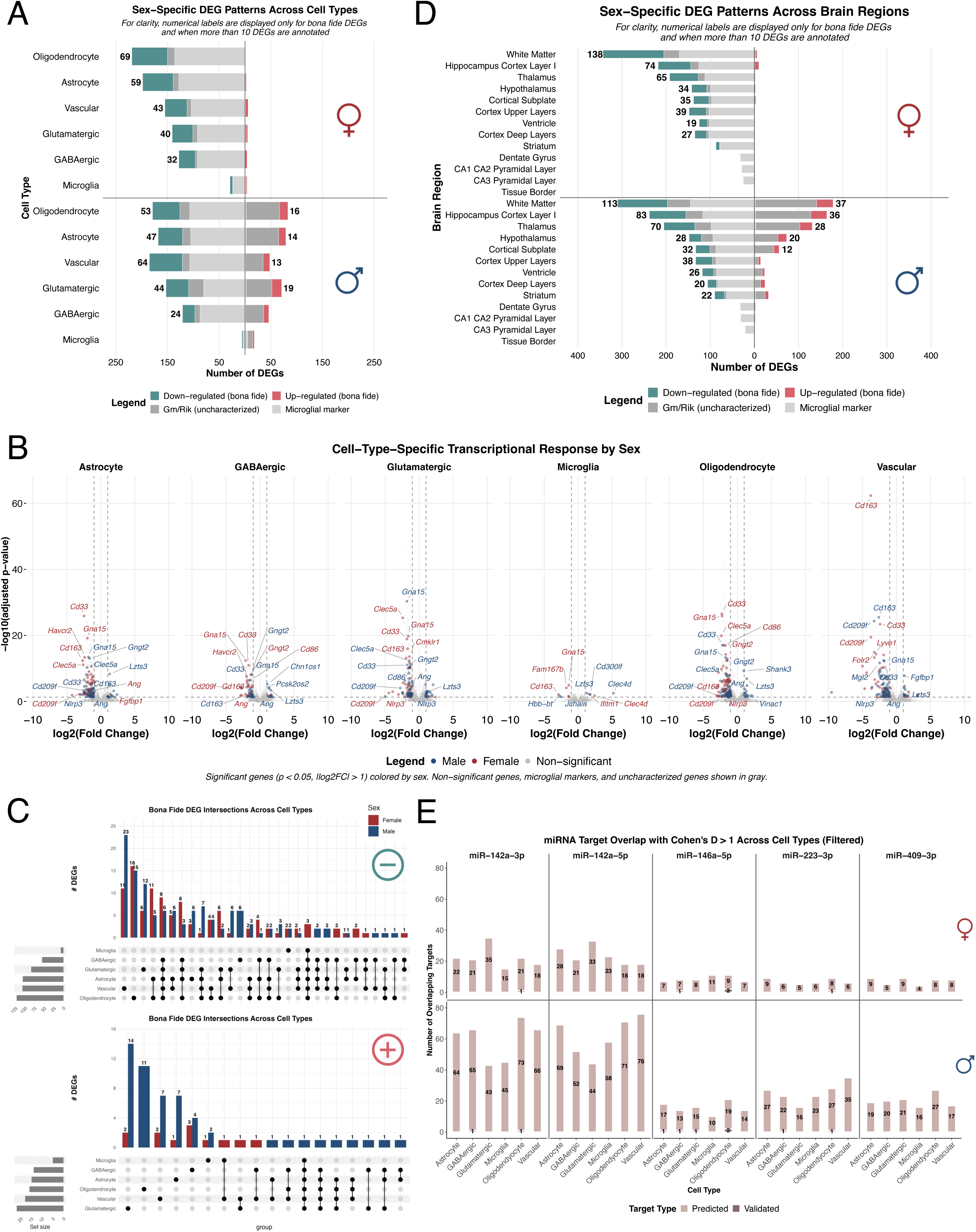
Microglia depletion triggers cell-type-specific transcriptional dysregulation with pronounced sex dimorphism and miRNA target enrichment. A. Bar plots showing a number of DEGs per cell type and sex from spatial transcriptomics. Bona fide DEGs are genes retained after excluding established microglial markers and genes lacking reliable cell-type annotation. Differential expression was determined by |log2FC| > 1.0, and adj p-value < 0.05. Females show predominantly downregulation across all cell types; males show bidirectional dysregulation, particularly in oligodendrocytes, astrocytes, and vascular cells. B. Volcano plots showing cell-type-specific transcriptional responses by sex. Points are color-coded by significance and sex: blue, significant DEGs in males; red, significant DEGs in females; gray, non-significant genes, microglial markers and uncharacterized Gm/Rik genes. The top 5 DEGs per sex (smallest p-value) are also labeled with gene names. Oligodendrocytes, astrocytes and vascular cells show the broadest effect size distributions in males, while females show more compressed distributions with higher statistical significance. C. UpSet plot showing overlap of DEGs across cell types for the top 10 intersections, separated by direction (downregulated: top panel; upregulated: bottom panel). Bar height indicates number of genes in each intersection; filled dots indicate which cell types participate. For downregulated genes, the largest intersections are predominantly cell-type-specific, with male vascular cells showing the largest single intersection (23 genes). For upregulated genes, the largest intersections are predominantly male-specific. D. Bar plots showing the number of DEGs per brain region and sex. Bona fide DEGs are genes retained after excluding established microglial markers and genes lacking reliable cell-type annotation. White matter-rich regions show the highest DEG counts, paralleling oligodendrocytes vulnerability at the cell-type level. E. Bar plots showing miRNA target enrichment of DEGs with Cohen’s D > 1.0 for the downregulated miRNAs. Target genes were identified using experimentally validated intersections from miRTarBase and predicted targets from TargetScanMouse. Bar heights indicate the number of overlapping targets (predicted + validated). Males show greater absolute numbers of miRNA target overlap across multiple cell types compared to females.

In females, oligodendrocytes showed the largest number of downregulated DEGs (69), followed by astrocytes (59), vascular cells (43), glutamatergic neurons (40) and GABAergic neurons (32). Consistent with the known CSF1R-dependence of border-associated macrophages (BAMs), which are molecularly distinct from parenchymal microglia^23^, spatially adjacent perivascular populations appear to contribute to transcriptional signal to neighboring cell-type bins. Notably, this signal was identified after applying the microglial marker exclusion filter. The most recurrent female downregulated signal, *Cd209a/b/d/f* and *Cd163*, across astrocyte, vascular, glutamatergic and GABAergic compartments (log2FC = -5.05 to -2.58; adj. p-value < 0.05), is consistent with co-capture of BAM-derived transcripts rather than intrinsic cell-type responses (expression profiles of text-referenced DEGs across cell types and experimental groups are shown in **Supp. Fig. 6**) (complete list of DEGs per cell type in **Supp. Table 4**).

Males exhibited a different pattern characterized by bidirectional dysregulation (**Fig. 3A**). While the DEGs cell-type hierarchy remained similar – vascular cells (64), oligodendrocytes (53), astrocytes (47) – males additionally exhibited pronounced upregulation across all cell types. Male oligodendrocytes had 16 genes upregulated versus no upregulated genes in females. The most consistently upregulated gene across all male cell types was *Lzts3* (log2FC = 1.2 to 1.4; adj. p-value < 0.01), which encodes a postsynaptic density scaffold that promotes dendritic spine maturation and binds directly to *Shank3*^32^. Among male oligodendrocytes, top upregulated genes included *Shank3* (log2FC = 1.1; adj. p-value < 0.001), and *Vinac1* (log2FC = 2.23; adj. p-value < 0.05) (**Supp. Fig. 6**, **Supp. Table 4**). While *Shank3* expression is canonically associated with neurons, its detection in oligodendrocytes is consistent with emerging evidence from non-neuronal roles in synaptic support and myelination-associated remodeling^33^. *Vinac1* currently lacks a characterized functional role in the brain. Male vascular cells displayed similar induction with *Fgfbp1* (log2FC = 1.07; adj. p-value < 0.001), alongside *Pcdha9* (log2FC = 2.96; adj. p-value < 0.05), and *Ccr7* (log2FC = 3.11; adj. p-value < 0.05). In male microglia, *Clec4d* was strongly upregulated (log2FC = 5.12; adj. p-value < 0.01). Beyond this upregulation, male cell types also showed recurrent downregulation of individual genes across multiple non-microglial populations. The most steadily suppressed gene was *Ang* (log2FC = -1.01 to -1.35; adj. p-value < 0.05), downregulated in astrocytes and GABAergic neurons of both sexes, and, in males, extending to glutamatergic neurons, oligodendrocytes and vascular cells. In females, *Ang* showed the same downward direction in glutamatergic neurons (adj. p-value < 0.05), but did not reach the fold-change threshold. *Nlrp3*, encoding the core inflammasome sensor, was downregulated in both sexes but with a different cell-type distribution: in glutamatergic neurons of both sexes, extending to astrocytes and vascular cells in males and to oligodendrocytes in females (log2FC = -2.47 to -1.25; adj. p-value < 0.05). It was non-significant in microglia of either sex. Although *Nlrp3* expression in the healthy brain is predominantly associated with microglia^34^, its presence in non-microglial cells remains an active area of investigation^35^, and its regulation in the rodent brain has been shown to be sexually dimorphic^36^. Whether the attenuation of this inflammatory priming reflects an adaptive response to microglial loss or a secondary consequence of disrupted microglia-to-cell signaling remains to be determined. Volcano plot visualization confirmed these sex- and cell-type-specific patterns (**Fig. 3B**). Oligodendrocytes, astrocytes, and vascular cells displayed the broad effect size distribution in males, with genes distributed across upregulated and downregulated states. Females showed more compressed distributions centered on downregulation.

Set analysis, visualized by UpSet plots, examined cross-cell-type sharing versus specificity of transcriptional responses by sex (**Fig. 3C**) (gene-level membership for all intersections shown in Fig. 3C is provided in **Supp. Table 5**). For downregulated genes, the two largest intersections were cell-type-specific: vascular cells showed the greatest suppressive response (11 female; 23 male), followed by oligodendrocyte-specific downregulation (16 female; 15 male), with smaller cell-type-specific intersections across the remaining populations. Multi-cell-type shared intersections were present but comprised smaller gene sets, and several included known myeloid marker genes (for example *Cd33, Cd86,* and *Clec5a*), likely reflecting spatial co-capture of PLX depleted BAMs. In contrast, for upregulated genes, the pattern was markedly male-biased. The two largest intersections were cell-type-specific and predominantly male: glutamatergic neurons (2 female; 14 male) and oligodendrocytes (11 male), followed by vascular (2 female; 7 male) and smaller contributions from other populations. Female-specific upregulation appeared only in small intersections throughout. This asymmetry indicates that gene induction following microglial depletion is stronger in males across multiple cell types, while suppressive responses show a more comparable sex distribution.

To examine whether these cell-type-specific transcriptional changes map onto anatomically defined structures, we performed a complementary analysis using brain region annotations (**Fig. 3D**; region definitions in **Supp. Fig. 7A-D**). White matter-enriched regions showed a higher number of DEG counts: 76 downregulated and 38 upregulated genes in males versus 92 downregulated and 6 upregulated genes in females. Regional boundaries remained consistent across all samples, enabling reliable pseudobulk comparisons (**Supp. Fig. 7A-D**). DEGs were calculated independently using pseudobulk aggregation within each anatomical region, providing a complementary spatial perspective to the cell-type analysis above. The predominance of downregulated genes in white matter across both sexes is consistent with the high microglial density in these regions, while the male-specific upregulation burden parallels the cell-type-level pattern described above.

To test whether DEGs were enriched for predicted and validated targets of downregulated miRNAs, we examined genes with strong effect sizes (Cohen’s D > 1.0) for targeting (**Fig. 3E**). Venn diagrams of genes passing the Cohen’s D > 1.0 threshold, stratified by cell type and sex, are provided in **Supp. Fig. 5D** (full list provided in **Supp. Table 6**). Across all miRNAs and cell types, males showed greater absolute target overlap than females (validated and predicted targets combined). The miR-142a arms accounted for largest target overlaps: in males, miR-142a-3p targets ranged from 43 (glutamatergic) to 74 (oligodendrocytes) genes per cell type, and miR-142a-5p from 44 (glutamatergic) to 76 (vascular), exceeding miR-146a-5p, miR-223-3p, and miR-409-3p (predicted and validated targets of each miRNA overlapping with DEGs in each cell type are listed in **Supp. Table 7**).

In females, validated targets included *Irak1* in oligodendrocytes, a miR-146a-5p target involved in TLR/NF-κB innate-immune signaling^37^. In males, the stronger enrichment extended to the miR-223-3p validated target *Igf1r* across GABAergic cells and the predicted synaptic-vesicle regulator *Vamp2*. To assess whether these targets converge on coherent functions rather than dispersed individual genes, we performed GO biological-process over-representation analysis on the male and female target sets. In males, the targets converged on synaptic and axonal organization, including axonogenesis, axon guidance and regulation of synaptic plasticity. Female target genes contained synaptic-annotated targets but showed no convergence surviving multiple-testing correction. These patterns are consistent with depletion-associated loss of microglia-enriched miRNAs co-occurring with elevated expression of their predicted and validated targets, a relationship more pronounced in males. While spatial transcriptomics revealed sex-specific transcriptional responses to microglial depletion, the functional consequences of these changes depend ultimately on protein-level alterations, which do not always track with mRNA abundance.

### Proteomic responses to microglial depletion are larger in males and independent from transcript-level changes

Bulk proteomics from the bregma -2.06 coronal block revealed sex-dimorphic protein changes. Quality control metrics confirmed consistent proteome coverage across all samples (4,848 proteins detected), with high inter-sample correlations (0.85-1.0) and comparable intensity distributions (**Supp. Fig. 8A-E**). Males exhibited 295 differentially expressed proteins (DEPs) compared to 34 in females – 8.7-fold difference exceeding the transcriptional sex dimorphism (**Fig. 4A-B, Supp. Table 8**).

**Figure 4:**
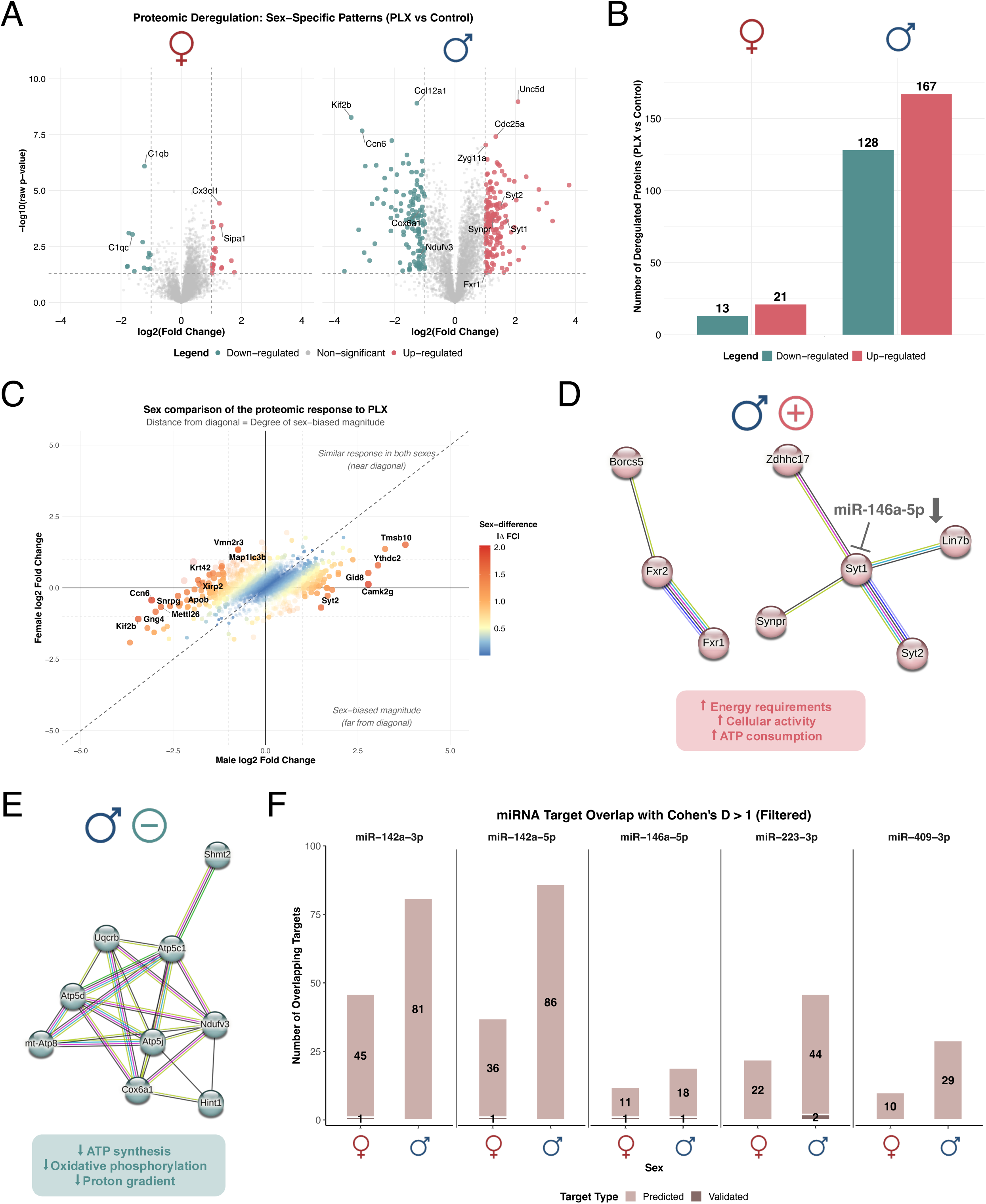
Sex-dimorphic proteomic responses to microglial depletion, independent from transcript-level changes. A. Volcano plots showing proteomic differential expression in females (left) and males (right) from bulk proteomics. Differential expression was determined by |log2FC| > 1.0, and raw p-value < 0.05. Males exhibit 295 DEPs versus 34 in females. Labeled proteins are those referenced in the main text. B. Stacked bar plot quantifying DEPs by sex. Males show 167 downregulated and 128 upregulated proteins; females show 13 downregulated and 21 upregulated proteins. C. Sex-comparison plot showing male log2FC (x-axis) versus female log2FC (y-axis) for all proteins. Diagonal dashed line indicates a similar response; distance from diagonal indicates sex-biased regulation. Point color represents the sex-difference score (|ΔFC| between sexes). Labeled proteins show high sex-specific responses. Most male DEPs show minimal changes in females. D. STRING protein-protein interaction network of upregulated proteins in males. Edge thickness indicates interaction confidence. Network includes synaptic vesicle protein (SYT1/2, SYNPR) and signaling regulators (FXR1/2). SYT1 is a predicted target of miR-146a-5p. E. STRING protein-protein interaction network of downregulated proteins in males. Network comprises mitochondrial ATP synthesis machinery including respiratory chain complexes (NDUFV3, COX6A1) and ATP synthase subunits (ATP5G/L). No significant networks identified in females. F. Bar plots showing miRNA target enrichment of DEPs with Cohen’s D > 1.0 for the downregulated miRNAs. Target proteins were identified using experimentally validated intersections from miRTarBase and predicted targets from TargetScanMouse. Bar heights indicate the number of overlapping targets (predicted + validated). Males show greater absolute numbers of miRNA target overlap compared to females.

Males exhibited 167 downregulated proteins, including COL12A1 (log2FC = -1.27; raw p-value < 0.001), KIF2B (log2FC = -3.44; raw p-value < 0.001), and CCN6 (log2FC = -3.08; raw p-value < 0.001). Conversely, 128 proteins were upregulated, including UNC5D (log2FC = 2.09; raw p-value < 0.001), CDC25A (log2FC = 1.35; raw p-value < 0.001), and ZYG11A (log2FC = 1.02; raw p-value < 0.001). In contrast, females had 13 downregulated proteins, primarily complement components C1QB (log2FC = -1.22; raw p-value < 0.001) and C1QC (log2FC = -1.62; raw p-value < 0.001), and 21 upregulated proteins, including CX3CL1 (log2FC = 1.26; raw p-value < 0.001) and SIPA1 (log2FC = 1.32; raw p-value < 0.001) (expression profiles of text-referenced DEPs across experimental groups are shown in **Supp. Fig. 9A-H**). Notably, the female downregulated set was dominated by complement components, consistent with direct loss of microglial protein products, whereas female dysregulation extended broadly across functional categories.

Comparison of male and female log2 fold changes revealed predominantly sex-biased proteomic responses (**Fig. 4C**). Proteins near the diagonal change to a similar extent in both sexes, whereas distance from the diagonal reflects the degree to which one sex responds more strongly (the quadrant indicates whether the two sexes change in the same or opposite direction). Proteins driving the off-diagonal spread, such as SYT2, TMSB10, and GID8, changed strongly in males but little in females. This suggests a male-biased response in magnitude rather than opposite-direction regulation. Together, these patterns indicate that microglial depletion engages a larger proteomic program in males than in females.

STRING protein-protein interaction analysis of all male DEPs revealed two distinct functional clusters (**Fig. 4D-E**). Upregulated proteins (**Fig. 4D**) clustered in pathways related to synaptic activity and vesicle trafficking, including synaptic vesicle proteins (SYT1/2, SYNPR) and signaling regulators (FXR1/2). Downregulated proteins (**Fig. 4E**) formed an interconnected network of mitochondrial energy metabolism components, including ATP synthase subunits (ATP5G/L) and respiratory chain complexes (NDUFV3, COX6A1). The co-occurrence suggests a potential imbalance between energy-consuming and energy-producing processes in males, though functional validation would be required to establish metabolic consequences.

To test whether proteomic dysregulation was associated with loss of microglia-enriched miRNAs, we examined proteins with strong effect sizes (Cohen’s D > 1.0) for overlap with targets of the downregulated miRNAs (**Fig. 4F**). Filtering for strong effect sizes retained 1735 proteins in males and 1070 in females, with 640 proteins shared between the two sexes (**Supp. Fig. 8F**; full list provided in **Supp. Table 6**). Similarly to spatial transcriptomics, the miR-142a arms showed the largest overlaps (validated and predicted targets combined): males showed 81 (miR-142a-3p) and 86 (miR-142a-5p) target proteins versus 46 (miR-142a-3p) and 37 (miR-142a-5p) in females, respectively. Across every miRNA examined, male overlap exceeded female: miR-223-3p showed 46 target proteins in males vs 22 in females, miR-146a-5p showed 19 vs 12, and miR-409-3p showed 29 vs 10. Cell-type-resolved spatial transcriptomics analysis revealed that these targets are expressed across multiple neuronal and glial populations. For example, synaptic genes such as presynaptic vesicle proteins *Syt1* and *Vamp2* and the postsynaptic density scaffold *Dlgap1* were enriched in neuronal populations. Synaptic vesicle and postsynaptic components of this kind are among the gene classes most consistently implicated in brain aging^38^.

Consistent with the well-documented divergence between steady-state mRNA and protein levels^39^, the protein-level dysregulation occurred without corresponding transcriptional changes. Despite the widespread expression, none of the deregulated protein-level miRNA targets showed corresponding significant mRNA changes in any cell type or sex. For example, SYT1, a predicted miR-146a-5p target, displayed particularly strong upregulation in males (log2FC = 1.68; raw p-value < 0.001), without corresponding mRNA changes (predicted and validated targets of each miRNA overlapping with DEGs and DEPs in each cell type and data layer are listed in **Supp. Table 7**). Targets of the downregulated miRNAs, that were strongly dysregulated at the protein level (Cohen’s D > 1.0), converged significantly on synaptic and axonal organization, but showed no comparable convergence in females.

Males experienced pronounced proteomic dysregulation following microglial loss, while females showed limited detectable proteomic change. The 8.7-fold greater proteomic dysregulation in males compared to females represents the largest relative sex difference across all three omic layers. In males, this pattern (increased synaptic proteins alongside decreased ATP-synthesis machinery) suggests a potential imbalance between energy-consuming and energy-producing processes, which was absent in females and would require additional functional validation to confirm.

## Discussion

This study reveals that aged male and female brains respond differently to microglial depletion. Males show more pronounced proteomic dysregulation, accompanied by directionally opposing shifts in synaptic and mitochondrial proteins, whereas females show comparatively limited proteomic change and no such pattern. These findings challenge assumptions of sexually monomorphic microglial function and have implications for therapeutic strategies.

While sex differences in microglial transcriptomes have been documented^40,41^, their functional consequences for neuronal homeostasis remained unclear. Recent work demonstrated that microglial response to pathology is sexually dimorphic in human post-mortem Alzheimer’s disease (AD) brain tissue^42^. Our depletion approach extends these observations by associating sex differences with distinct molecular signatures rather than phenotypic variation alone.

The near-complete depletion of microglia-enriched miRNAs across brain regions indicates that a large fraction of the bulk signal for these miRNAs is derived from microglia. Whether microglia are a quantitatively substantial source of regulatory small RNAs for other cell types cannot be resolved from bulk data, but is plausible given prior evidence that microglia release extracellular vesicles containing functional miRNAs that transfer to neurons and modulate translation^43,44^. These miRNAs have validated targets in inflammatory and translational control pathways: miR-142a targets SOCS1/TGFBR1^45^, miR-146a-5p suppresses IRAK1/TRAF6^46^, and miR-223-3p inhibits NLRP3 inflammasome^47^. Comparable miRNA target enrichment among high-effect-size proteins in both sexes, with a modest male-predominant trend, suggests that aged male brains may operate closer to a regulatory threshold, such that loss of microglial input is more readily reflected at the protein level. The smaller proteomic responses in females, despite similar enrichment proportions, suggest tighter basal transcriptional control or alternative compensatory mechanisms that buffer against microglia loss. We emphasize this as a hypothesis with the observed overlap rather than a demonstrated dependency, as our design tests neither causality nor miRNA transfer directly.

Males show 8.7-fold more DEPs than females despite modest transcript differences. Similar transcriptome-proteome discordance characterizes neurodegenerative diseases: only 37% of proteins correlated with mRNA in AD dementia^48^. The concurrent increase in synaptic proteins and decrease in mitochondrial ATP-synthesis components in males describes a potential imbalance between energy demand and supply at the protein level. Given that microglia can transfer mitochondria to stressed neurons^49^, this male-specific pattern potentially suggests that aged male neurons depend on microglial metabolic support, though whether it reflects a functionally consequential state remains to be determined.

These findings may help explain inconsistent results in CSF1R inhibitor trials. Pexidartinib, the most clinically advanced CSF1R inhibitor, has shown variable efficacy across settings despite consistent target engagement^50^. While it received FDA approval for tenosynovial giant cell tumor, it carries a black box warning for hepatotoxicity^51^, and microglial depletion in glioblastoma yielded no clinical improvement^52^. Sex-specific responses have also been reported: only females showed functional rescue and extended survival in tauopathy models despite similar depletion^53^, while males exhibited greater depletion efficacy^19^.

Several limitations warrant consideration. Three-week PLX treatment represents acute depletion rather than the chronic microglial loss of natural aging or disease. While long-term CSF1R inhibition can achieve >95% depletion for up to 6 months^13^, our protocol may trigger compensatory responses distinct from gradual dysfunction. The approximately 75% depletion achieved leaves surviving microglia that could represent a specialized subpopulation with distinct functions. The study examines a single aged timepoint; single-cell analyses reveal cell-type-specific aging trajectories^54^, suggesting that sex-dimorphic microglial dependencies may emerge at specific life stages and may not generalize across the full aging continuum. Given the exploratory nature of this study, we used raw p-values for primary differential expression analysis of miRNAs, with adjusted p-value results reported alongside in supplementary materials.

PLX5622 is not fully microglia-selective and can affect peripheral myeloid populations including circulating monocytes. Excluding a peripheral contribution would require flow cytometric profiling of circulating monocytes, which was not performed here and would need a newly generated cohort to address this. However, our compositional analysis detected no significant change in vascular-associated cells following PLX treatment, the compartment through which peripheral monocytes would enter the parenchyma.

Female control samples exhibited some biological variance. This variability likely reflects the biology of aged female mice, which at 21 months are in a post-reproductive phase of asynchronous reproductive senescence, producing heterogenous hormonal states ranging from intermittent cycling to full acyclicity^55^. By this age ovarian reserve, estradiol and inflammatory cytokine profiles diverge substantially across individuals^56^. Compounded by the well-established increase in transcriptional noise with aging^57,58^, this variability is specific to the aged context. Importantly, young adult female rodents are not inherently more variable than males^59,60^, so heterogeneity observed here potentially reflects genuine biology rather than a systemic sex confound. The partial clustering of some female control with PLX-treated samples may reflect this, though contributions from technical sources cannot be excluded. We therefore interpret female-specific findings conservatively. Similarly, the pronounced proteomic dysregulation observed in males should be interpreted with awareness that small sample sizes limit effect-size precision. Replication in larger independent cohorts will be necessary to confirm the degree of sex dimorphism reported here.

Finally, spatial transcriptomics captures mRNA at cell locations, but transcript diffusion may blur cell-type-specific signals, particularly in high cellular density regions or where cell boundaries are difficult to resolve. To address this, consensus downregulated genes appearing across multiple cell types were cross-referenced against a microglial reference expression atlas, confirming predominant identity as microglial markers rather than cell-type-specific transcriptional responses.

In summary, aged male and female brains respond differently to microglial depletion, revealing sex-dimorphic patterns of molecular vulnerability. The data show a larger proteomic male response, and this pattern is suggestive of sex-specific compensatory mechanisms, though the precise nature of these mechanisms remains to be established. These results do not argue against microglial depletion as a therapeutic strategy but emphasize the need for sex-specific approaches. Precision medicine approaches accounting for sex, age, and metabolic status may prove important for successful microglial-targeting therapies.

## Methods

### Animal procedures

Male and female C57BL/6J mice were obtained from National Institute of Health – National Institute of Aging colony. Mice were housed under a 12h light/dark cycle and provided with water and standard chow ad libitum. All animal care and procedures were conducted in accordance with institutional guidelines approved by Administrative Panel on Laboratory Animal Care at Stanford University. For microglial depletion, mice received PLX5622 produced as a 1200mg drug per 1kg diet (1200ppm) for three consecutive weeks. Control mice received open standard diet with 15kcal% fat.

Following treatment, mice were anesthetized with isoflurane and transcardially perfused with PBS. Brains were dissected, kept whole, and immediately snap frozen for approximately 20 seconds in liquid nitrogen-cooled isopentane. Samples were stored at -80□C until further processing. Cryopreserved brain tissue was embedded in Tissue Tec OCT embedding medium (Leica; Wetzlar, Germany). Each brain was cut into 10□m slices, which were further used for downstream experiments. No animals were excluded from any omics analysis.

### Immunofluorescence labeling of mouse brain tissue

Brain tissue was postfixed with 4% paraformaldehyde (PFA) for 24 h directly after tissue extraction, cryoprotected in 30% sucrose for 3 nights and subsequently embedded in Tissue-Tek OCT cryo-embedding matrix prior to freezing the tissue on dry ice. Brain samples were cut into 16µm sections (CM3050 S cryostat, Leica) and stored at -80°C. Immunofluorescence staining was performed on sections that were adjusted to RT for 30 min and washed with PBS to remove excess embedding matrix. Sections were transferred to citrate buffer (0.01 M, pH 6.0) for antigen retrieval. Tissue permeabilization was performed by incubating sections with 0.3% Triton-X-100 in PBS for 10 min. Blocking was performed using a mixed blocking solution (2.5% bovine serum albumin, 2.5% fish gelatin, 2.5% fetal calf serum in PBS). The primary antibodies (Iba1 (1:250, abcam, ab5076), TMEM119 (1:250, abcam, ab210405), CD31 (1:50, Dianova, DIA-310), CD206 (Cell Signaling Technology, 24595T)) were diluted in antibody diluent (0.625% bovine serum albumin, 0.625% fish gelatin, 0.625% fetal calf serum in PBS) and incubated for 24h at 4°C, followed by incubation with species-specific Alexa Fluor-coupled secondary antibodies (1:500, Thermo Fisher Scientific), for 2h at RT. Afterwards, sections were incubated with DAPI (1:1000, Thermo Fisher Scientific, 62248) for 10min and mounted with ProLong™ Gold Antifade mounting solution (Invitrogen, P36930).

#### Image acquisition

Sagittal sections were imaged at 10x and representative detailed views at 40x using a Zeiss LSM 900 confocal microscope with a 10x (PlnApo10x/0.45 DICII) and a 40x (PlnApo 40x/1.4 Oil DICII) objective. 10x image mosaics were merged and 40x image stacks were processed using the Zeiss Zen Blue 3.2 software suite.

### MiRNA sequencing

#### Sample preparation

RNA was isolated from 5-10 individual 10□m sections per region using the miRNeasy Mini Kit (Qiagen; Hilden, Germany); for the bregma -2.06 region, sections were cut immediately anterior to the section used for spatial transcriptomics. RNA integrity was assessed using the RNA 6000 Nano Bioanalyzer Kit (Agilent Technologies; Santa Clara, CA, USA), yielding RNA integrity numbers (RIN) > 7.5 for all samples (average RIN value of 8.6, range 7.9 – 9.5) (per-sample values in **Supp. Table 9**). Small RNA library preparation was performed using the MGIEasy Small RNA Library Prep Kit (Cat. No. 940-000196-00; MGI Tech, Shenzhen, China) on the high-throughput MGI SP960 automated sample prep system following the manufacturer’s instructions. Briefly, 3’- and 5’-adapters were ligated to the RNA, followed by reverse transcription (RT) with an RT primer containing sample-specific barcodes. The resulting cDNA was amplified by PCR, size-selected and purified using AMPure XP Beads (Beckman Coulter). Library quality was verified using an Agilent DNA 1000 kit (Agilent Technologies), and concentration was determined by Qubit 1X dsDNA High Sensitivity assay (Thermo Fisher Scientific). For each library, 18-20 barcoded samples were combined in equimolar ratios. Pooled libraries were circularized and sequenced on a DNBSEQ-G400RS instrument at the NGS Sequencing Facility of Saarland University using 50bp single-end sequencing.

#### Data analysis, statistics and reproducibility

Sequencing data were processed using miRMaster 2.0^61^ with the following parameters. 3’ adapter sequences were detected using a sliding window of 10 nucleotides (nt) requiring a minimum overlap of 4nt and a maximum edit distance of 1nt. A minimum PHRED quality score of 20 was required within the sliding window. Samples were excluded if fewer than 1 million reads were detected. Retained reads were aligned to the mouse GRCm38 primary assembly using Bowtie, allowing one mismatch per read. MiRNAs were annotated using miRBase v22.1^62^; other non-coding RNAs were quantified using GtRNAdb v18.1^63^, Ensembl ncRNA v100^64^, and NONCODE v5^65^. To ensure robust detection, only miRNAs detected at threshold of 5 raw counts, in at least 75% of samples, were included in downstream analyses.

Differential expression analysis was performed using Student’s t-test and independently for each brain region and sex, comparing PLX-treated to matched control samples. Given the exploratory aim of identifying candidate microglia-associated miRNAs for downstream analyses, miRNAs were primarily defined as using raw p-value < 0.05 and |log2FC| > 1.0. Benjamini-Hochberg (BH) adjusted p-values applying the same cut offs as before are reported in parallel as a sensitivity analysis. Downstream analyses were conducted in RStudio (v4.3.3).

### Spatial transcriptomics

#### Sample preparation

Cryopreserved tissues meeting the BGI Stereo-seq Transcriptomics T kit v1.3 (BGI; Shenzhen, China) protocol minimum of RIN >= 4 were used for spatial transcriptomics using the Stereo-seq Transcriptomics T kit v1.3 (BGI; Shenzhen, China) according to the manufacturers protocol. All samples had RIN values between 7.9 and 9.5, with an average RIN of 8.6 (per-sample values in **Supp. Table 9**). Briefly, a 10□m tissue sections were placed on the Stereo-seq chip, incubated at 37□C for 5min and fixed in ice-cold methanol at -20□C for 30min. Subsequently, nuclei were stained with fluorescent DNA dye (Qubit ssDNA kit; Thermo Fisher Scientific) and imaged on a MOTIC microscope (Hong Kong, China) in FITC channel to generate an image containing the spatial information of the cells. Tissues were permeabilized for 18min to release mRNA for capture by the spatially barcoded DNBs on the chip. Captured mRNA was reverse transcribed at 45□C for 2h.

After reverse transcription, remaining tissue was removed, generated cDNA was collected and amplified according to the manufacturers protocol. For each sample, 100ng of cDNA was used for library construction and DNB generation using the Stereo-seq 16 Barcode Library Preparation kit. Libraries were sequenced on the DNBSEQ-T10 sequencing platform (MGI; Riga, Latvia).

#### Data analysis, statistics and reproducibility

Sequenced libraries were processed using the SAW pipeline (https://github.com/STOmics/SAW). For read 1, coordinate identity (CID) sequences were mapped to the predefined coordinates of the in situ capture chip, allowing one mismatch. Unique molecular identifiers (UMIs) containing ambiguous bases (N) or more than two bases with quality scores <10 were removed. The corresponding CID and UMI information extracted from read 1 was appended to the header of the paired read 2. Retained read 2 sequenced were aligned to the mouse reference genome (mm10) using STAR^66^, and alignments with MAPQ >10 were kept. UMIs sharing the same CID and gene locus were collapsed, allowing one mismatch to account for sequencing errors.

SAW was used to generate spatial gene expression matrices at two resolutions: bin100 for region-level analyses and CellBin (image-guided cell segmentation from ssDNA staining) for cell-type-level analyses. Quality control (QC) analysis confirmed high technical quality across all samples. QC metrics, including total UMI counts, number of detected genes, and mitochondrial gene fraction, were computed for all cells. CellBin profiles were further refined by retaining only cells meeting quality thresholds of ≥200 genes, ≥300 UMIs and ≤15% mitochondrial content (**Supp. Fig. 10A**). QC filtering retained >97% of cells across all conditions without altering cell type composition (**Supp. Fig. 10B-C**), demonstrating that the vast majority of cell met high-quality standards. Inter-sample correlations within cell types ranged from 0.85-1.0 with microglia showing slightly lower correlations likely reflecting depletion-induced signal reduction (**Supp. Fig. 11A**).

For bin100 data, counts were normalized by library size (scaled to 10,000 counts per bin), and log-transformed. Highly variable genes were selected using a dispersion-based approach (min_mean = 0.0125; max_mean = 3; min_disp = 0.5), followed by scaling and PCA. To mitigate potential batch effects across samples, Harmony^67^ was applied in PCA space using sample identity as the batch variable. Leiden clustering was performed on Harmony-corrected embeddings. Anatomical regions were annotated based on anatomical landmarks identifiable from tissue morphology and their relative positions within each coronal section. For CellBin data, the top 3,000 variable genes were selected, and cell types were assigned based on canonical markers following standard normalization, PCA and Leiden clustering without batch correction.

Sex-stratified pseudobulk differential expression was performed using DESeq2^68^. Raw UMI counts were aggregated by summing gene-wise counts within each biological replicate for each anatomical region and cell type separately. Genes with total pseudobulk counts <10 were filtered prior to testing. To remove putative spatial co-capture artifacts from all cell types DEG lists, genes corresponding to established microglial markers were identified from three independent reference databases and excluded prior to downstream interpretation. Mouse microglia markers were retrieved from CellMarker 2.0^69^ and PanglaoDB^70^ (version March 2020), yielding 773 and 79 unique mouse microglia markers, respectively. A third set was obtained from the MSigDB M8 cell type signature collection^71^, restricted to two aged brain myeloid signatures, contributing 353 markers. In total, 1040 unique microglia markers were excluded for downstream analyses. Additionally, genes matching uncharacterized or unannotated patterns (Gm-prefixed and Rik-suffix genes) were excluded, as these lack functional annotation. DEGs were defined as adj. p-value < 0.05 and |log2FC| > 1.0. Genes highlighted in the Results and displayed in **Supp. Fig. 6** were selected using two criteria: recurrence, defined as reaching significance in >=3 cell types of one sex in the same direction, and cell-type-specific induction strength, defined as the largest log2FC among significant DEGs. Analyses were performed in Python (v3.8.20) using scanpy (v1.9.6), anndata (v0.8.0), numpy (v1.23.5), pandas (v1.5.3), scipy (v1.10.1), matplotlib (v3.7.5), pydeseq2 (v0.4.4) and RStudio (v4.4.3).

### MiRNA target database retrieval

Predicted miRNA targets were retrieved from TargetScanMouse (using Conserved Site Context Scores; accessed November 2025), retaining sites with a weighted context++ score percentile >=75 and deduplicated to one entry per gene symbol. Experimentally validated targets were obtained from miRTarBase (accessed November 2025) by combining entries from strong, weak and functional evidence files and deduplicating on target gene symbol. The two databases were used as parallel, independent lines of evidence rather than intersected with each other. To identify proteins and transcripts showing large-magnitude responses to microglial depletion, effect sizes were computed as Cohen’s D, calculated as the mean difference between PLX and control groups divided by the pooled standard deviation. Genes and proteins exceeding the effect-size threshold Cohen’s D > 1.0 were used for miRNA target analysis (defined in **Supp. Table 6**); full input target lists for both databases are provided in **Supp. Table 10**. Each target list was then separately intersected with the high-effect-size proteins or genes, and overlap counts are reported per database.

### Label-free quantitative proteomics

#### Sample preparation

Sample preparation was adapted from a previously established protocol^72^. Protein extraction was performed on 5-10 individual 10□m coronal sections cut immediately posterior to the bregma -2.06 section used for spatial transcriptomics. These were thawed at room temperature and resuspended in 0.4% SDS in PBS. Samples were lysed by sonication and protein extraction was determined by bicinchoninic acid (BCA) assay (Thermo Fisher Scientific). For each sample, the proteome amount was adjusted to 100□g.

Samples were precipitated overnight using ice-cold acetone, followed by centrifugation and an additional wash using ice-cold methanol. Protein pellets were resuspended in X-buffer (7M urea, 2M thiourea, and 20mM HEPES; pH 7.5) and reduced with dithiothreitol (DTT) (45min, 25□C, 450rpm). Cysteine residues were alkylated with iodoacetamide (IAA) in dark (30min, 25□C, 450rpm), and its surplus was quenched by adding DTT (45min, 25□C, 450rpm). Proteins were digested overnight with sequencing-grade trypsin (0.5□g/□L in 50mM acetic acid; Promega) (>16h, 37□C, 500rpm). Digestion was quenched with formic acid (FA), and peptides were desalted using 50mg C18 SepPak columns (Waters). Columns were equilibrated with acetonitrile (ACN), elution buffer (80% ACN / 0.5% FA), and 0.1% trifluoroacetic acid (TFA). Samples were loaded, washed with 0.1% TFA and 0.5% FA, and eluted with elution buffer under vacuum. Eluates were vacuum-dried (45□C) overnight, reconstituted in 1% FA with sonication, and filtered through 0.22□m centrifugal filters (Merck Millipore) before LC-MS/MS analysis.

Sample analysis was performed using a NanoElute nano flow LC system (Bruker, Germany) coupled to a timsTOF Pro mass spectrometer (Bruker, Germany) with a CaptiveSpray nanoESI source. Peptides were first loaded on a 5mm trap columns (Thermo Trap Cartridge) and washed with 6□L 0.1% FA with a flow rate of 10□L/min. Separation was performed on an Aurora Ultimate CSI analytical column (25cm x 75□m ID, 1.6□m FSC C18; IonOpticks) using a multistep gradient from 0% to 85% ACN in 0.1% FA over 100min at a flow rate of 400nL/min. Captive Spray nanoESI source (Bruker, Germany) was used to ionize the peptides at 1.5 kV with 180°C dry temperature at 3 L/min gas flow. The timsTOF Pro was operated in default dia-PASEF (data-independent acquisition parallel accumulation serial fragmentation) long-gradient mode. Trapped Ion Mobility Spectrometry (TIMS) settings were set to 1/K₀ start 0.6 Vs/cm² and end 1.6 Vs/cm², with ramp and accumulation times of 100ms each at 9.43 Hz. Mass range was set from 100.0 – 1700.0 Da (with positive ion polarity), with dia-PASEF range set to 400.0 – 1201.0 Da and mobility range 0.60 – 1.43 1/K₀ (cycle time of 1.80s). Collision energies were set to 20.0 eV for 0.60 1/K₀ and to 59.0 eV for 1.60 1/K₀. ESI Low Concentration Tuning Mix was used for calibration of m/z and mobility. The mass spectrometry proteomics data have been deposited to the ProteomeXchange Consortium via the PRIDE^73^ partner repository with the dataset identifier PXD074966.

#### Data analysis, statistics and reproducibility

Raw data were pre-processed using DIA-NN (v1.8.1)^74^ with the mouse UniProt reference proteome (downloaded August 2025) and a library-free search strategy. The precursor charge range was set to 2-4, with carbamidomethylation of cysteine as a fixed modification. Cross-run normalization was performed using DIA-NN’s RT-dependent normalization mode, and protein-level quantification was computed using DIA-NN’s implementation of the MaxLFQ algorithm^75^. Subsequent analysis was performed in Perseus (v2.1.5.0)^76^ using the resulting MaxLFQ-normalized protein-group matrix; no additional normalization was applied in Perseus. Intensity values were log2-transformed and filtered to retain proteins with at least three valid values per experimental group. Missing values were imputed per sample column using a normal distribution. with width 0.3 and downshift 1.8 (Perseus default parameters). This imputation strategy was selected because log2-transformed protein intensities are approximately normally distributed^77^, so imputed values are statistically consistent with the observed data. Furthermore, this choice is supported by benchmarking studies recommending normal distribution imputation for left-censored missing-not-at-random (MNAR) data in label-free quantitative proteomics^78,79^.

Sex-stratified differential expression analysis was performed using two-sample Student’s t-tests (unpaired, two-sided) comparing PLX-treated versus control groups. Statistical significance was assessed using permutation-based false discovery rate (FDR). DEPs were defined as p-value < 0.05 and |log2FC| > 1.0. Proteins highlighted in the Results and displayed in **Supp. Fig. 9A-H** were selected using two criteria: statistical rank, defined as DEPs with the smallest raw p-value per sex, and network membership, defined as proteins forming the STRING-based PPI modules discussed in the Results. Analyses were performed in RStudio (v4.4.3).

## Supporting information

Cell type proportion statistics comparing PLX-treated and control mice, stratified by sex.

Differential expression analysis of detected miRNAs following microglial depletion across brain regions, stratified by sex.

Cell-type enrichment scores for PLX-deregulated miRNAs from Pomper et al.

DEGs per cell type and sex following microglia depletion.

Cross-cell-type intersections of bona fide DEGs following microglia depletion.

miRNA target genes overlapping with DEGs and DEPs.

DEPs identified by proteomics following microglia depletion.

RNA integrity numbers (RIN) for all samples.

Predicted and validated target gene lists for candidate miRNAs.

## Data availability

The sequencing data generated in this study have been deposited in the NCBI’s Gene Expression Omnibus database under accession codes **GSE323GG3** and **GSE326105** for the miRNA-seq and spatial transcriptomics, respectively. The mass spectrometry proteomics data have been deposited to the ProteomeXchange Consortium via the PRIDE^73^ partner repository with the dataset identifier **PXD074G66**.

## Code availability

Custom analysis scripts used in this work are available from GitHub at https://github.com/amilabeg97/plx_microglia_depletion.

## Acknowledgements and funding

We thank all members of Keller and Wyss-Coray lab for feedback and support. This study is funded by the M.J. Fox Foundation (MJFF-021418; A.K. and T.W-C.), ASAP Collaborative Research Network (ASAP-027071; A.K.), Hans und Ruth Giessen Stiftung (V.W.), NIH Pathway to Independence Award (1K99AG088304-01, I.H.G.), MAC3 Dementia and Ageing Fellowship (I.H.G.).

## Author contributions

Animal treatments and brain tissue collection were performed by I.H.G. Bulk sequencing and spatial transcriptomics experiments were performed by A.B., N.L., and Va.W. Immunohistochemistry was performed by M.R., E.D.D. and Vi.W. Proteomics experiments were performed by A.B. and P.D., with supervision by R.M. Data analysis was led by A.B., with support from M.F., J.L., S.G., P.D. and S.R. Data interpretation was supported by A.K., Vi.W. and M.F. The manuscript was written by A.B. and reviewed by A.K., Vi.W., T.I. and M.F. The project was supervised by A.K., Vi.W. and T.W-C. Funding was acquired by A.K., Vi.W. and T.W-C. All authors read and approved the final manuscript.

## Ethics declarations (Competing interests)

A.K. is a member of the scientific advisory board of Firalis. The remaining authors declare no competing interests.

---

**Supplementary figure 1:**
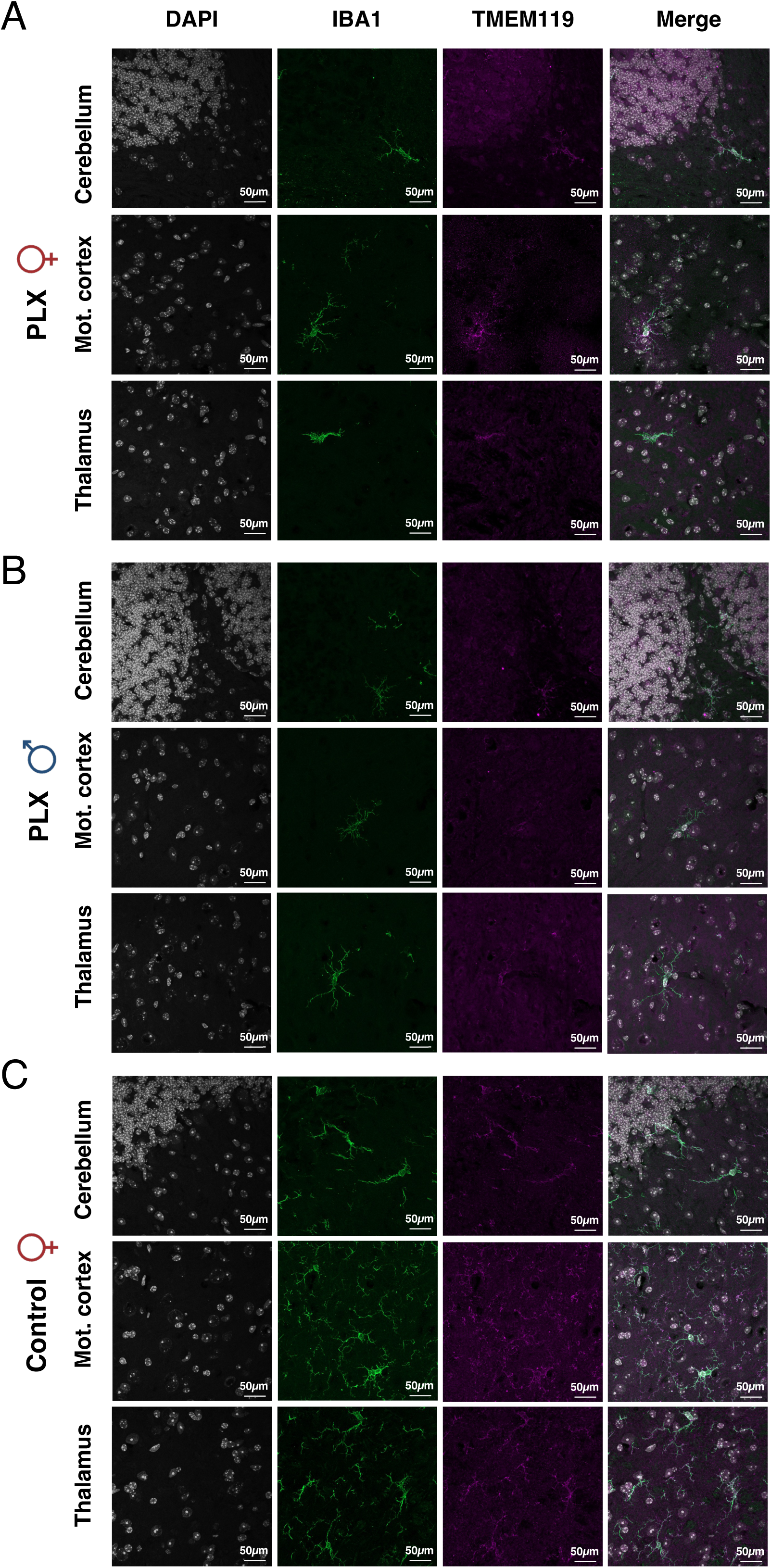
Immunofluorescence validation of microglial depletion in the brain tissue from 21-month-old mice using parenchymal microglial markers. Representative confocal images (40x magnification) of cerebellum, motor cortex, and thalamus (rows) stained for DAPI (gray), IBA1 (green), TMEM119 (magenta), shown as individual channels and merged. (A) PLX-treated female. (B) PLX-treated male. (C) Control female.

**Supplementary figure 2:**
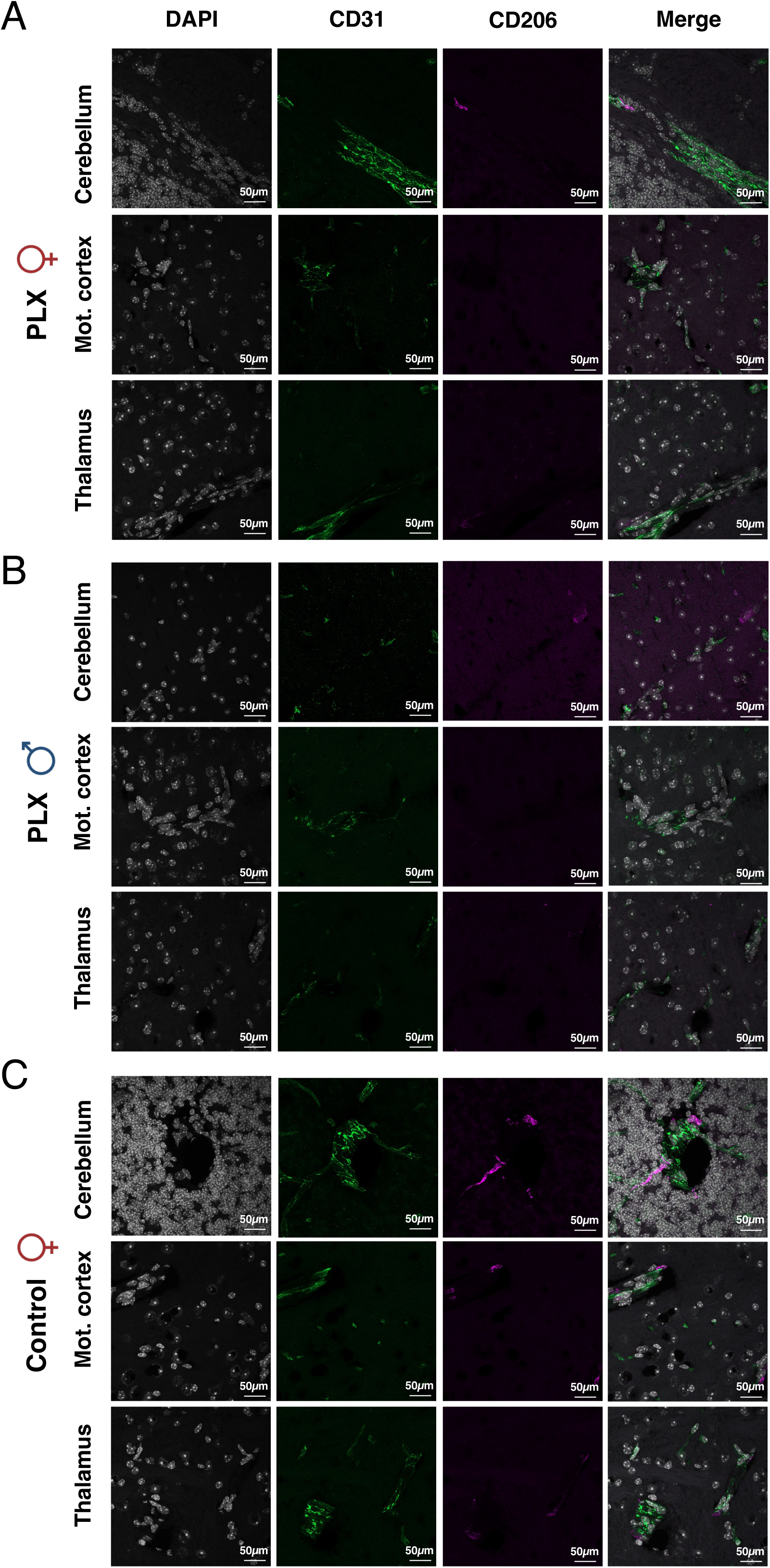
Immunofluorescence assessment of border-associated macrophage and vascular marker distribution following microglial depletion in 21-month-old mice. Representative confocal images (40x magnification) of cerebellum, motor cortex, and thalamus (rows) stained for DAPI (gray), CD31 (green), CD206 (magenta), shown as individual channels and merged. (A) PLX-treated female. (B) PLX-treated male. (C) Control female.

**Supplementary figure 3:**
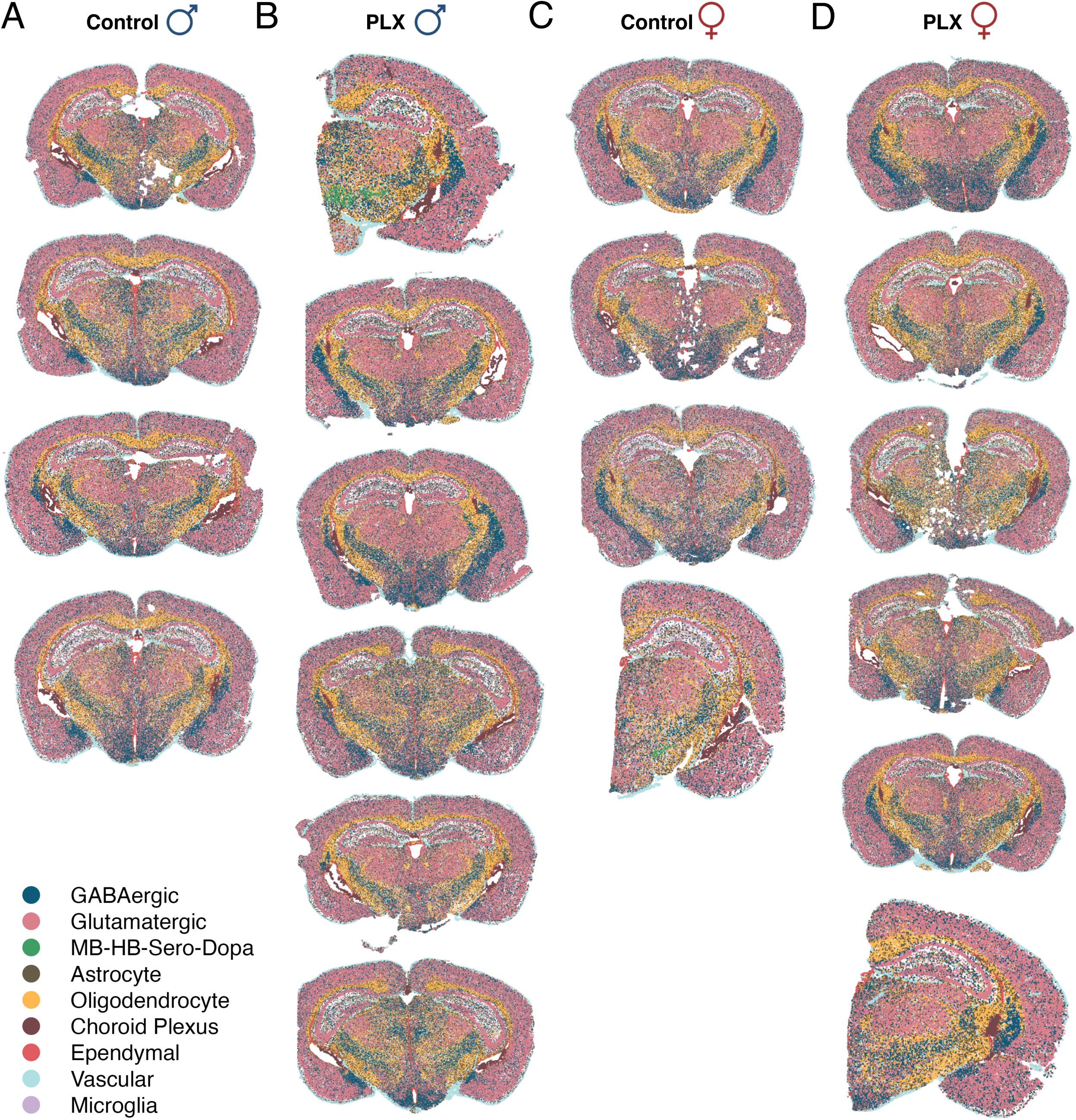
Spatial cell type distribution across experimental conditions. Spatial maps showing the distribution of six major annotated cell types across all samples. Each spot is colored by the cell type identity (legend in the bottom left corner). A. Control male mice (n = 4). B. PLX-treated male mice (n = 6). C. Control female mice (n = 4). D. PLX-treated female mice (n = 6).

**Supplementary figure 4:**
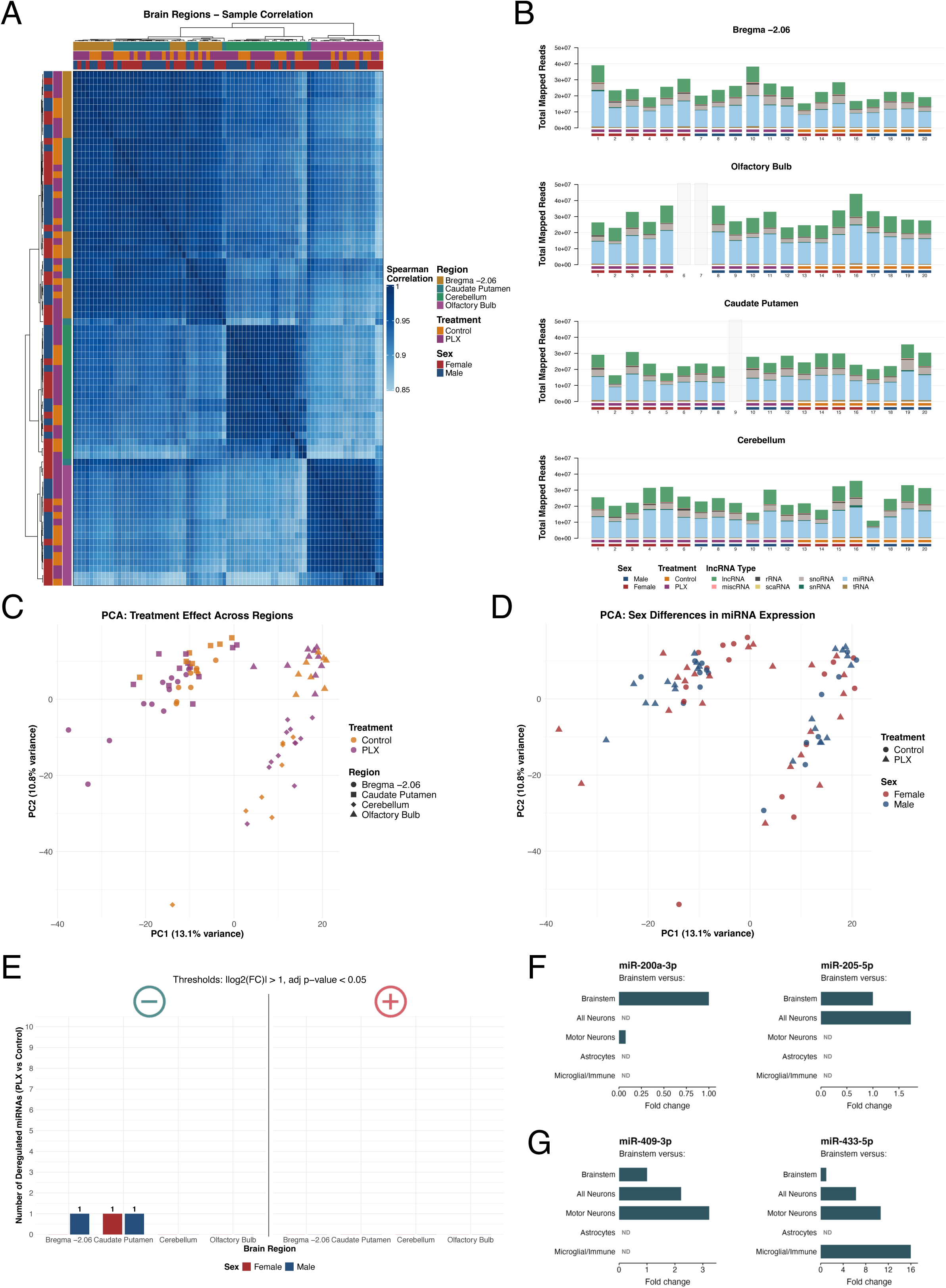
Quality control metrics and regional distribution of small RNA sequencing data. A. Heatmap showing Spearman correlation coefficients between all samples across four brain regions, treatment and sex. Samples clustered primarily by brain region rather than treatment or sex, indicating regional identity is the dominant source of variance in miRNA expression. Color bars indicate brain region, treatment condition and sex. B. Stacked bar plot showing total mapped reads per sample separated by the type of non-coding RNA (ncRNA) for each brain regions. MiRNA represent the dominant fraction of mapped reads across all samples. Bar height represents the number of total mapped reads, indicating consistent library quality and composition across experimental groups. Gray-shaded bars indicate samples not collected for that region. C. Principal component analysis (PCA) of miRNA expression patterns showing treatment effect across brain regions. Shape indicates brain region; color indicates treatment. PC1 primarily separates samples by anatomical region. The overlapping distribution of Control and PLX samples within each region indicates that regional variation exceeds treatment-induced changes in global miRNA expression profiles. D. PCA showing sex-specific differences in miRNA expression patterns. Shape indicates treatment; color indicates sex. The relatively overlapping distribution of male and female samples suggests that sex differences explain less variance than regional variation in miRNA expression profiles. E. Stacked bar plot showing the number of deregulated miRNAs per brain region and sex using the FDR-adjusted p-values. Differential expression was determined by |log2FC| > 1.0, and adjusted p-value < 0.05. Numbers indicate count of downregulated and upregulated miRNAs (colors indicate sex). F. Fold change in expression of miR-200a-3p and miR-205-5p across CNS populations (neurons, motor neurons, astrocytes, microglia/immune cells) relative to brainstem tissue, derived from Pomper et al.^24,30^ CNS miRNA expression database. G. Same-cell-type fold change comparison as in F, but for miR-409-3p and miR-433-5p.

**Supplementary figure 5:**
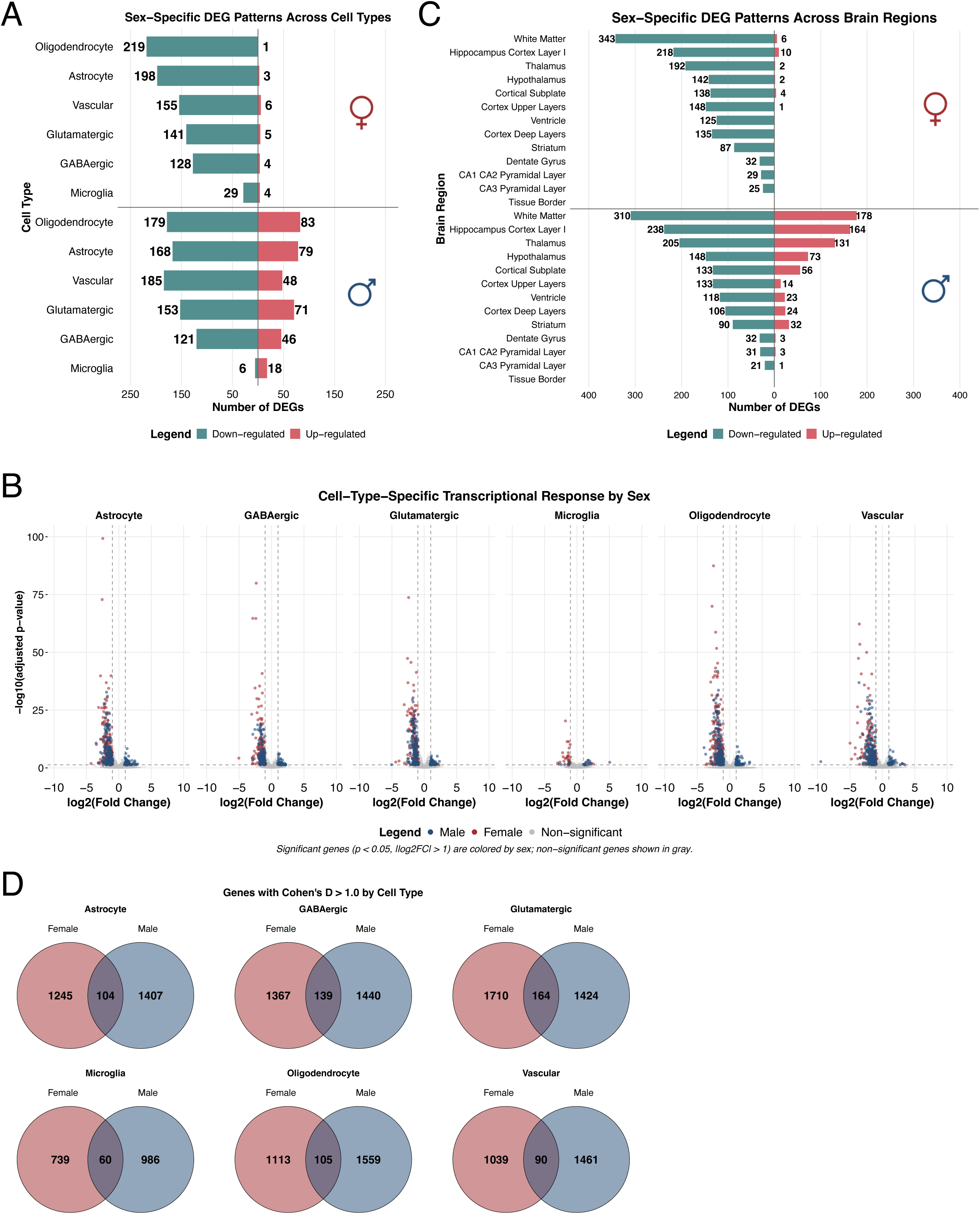
Cell-type specific DEG counts prior to microglial marker gene and Gm/Rik gene exclusion. A. Bar plot showing a number of DEGs per cell type and sex from spatial transcriptomics. Differential expression was determined by |log2FC| > 1.0, and adjusted p-value < 0.05. B. Volcano plots showing cell-type-specific transcriptional responses by sex. Blue points: significant in males; purple: significant in females; gray points: non-significant. C. Bar plot showing the number of DEGs per brain region and sex. White matter-rich regions show the highest DEG counts, paralleling oligodendrocytes vulnerability at the cell-type level. D. Venn diagrams showing the number of genes with Cohen’s D > 1.0, per cell type, with the overlap between sexes. These effect-size-filtered gene sets served as input for the miRNA target-overlap analysis in Fig. 3E.

**Supplementary figure 6:**
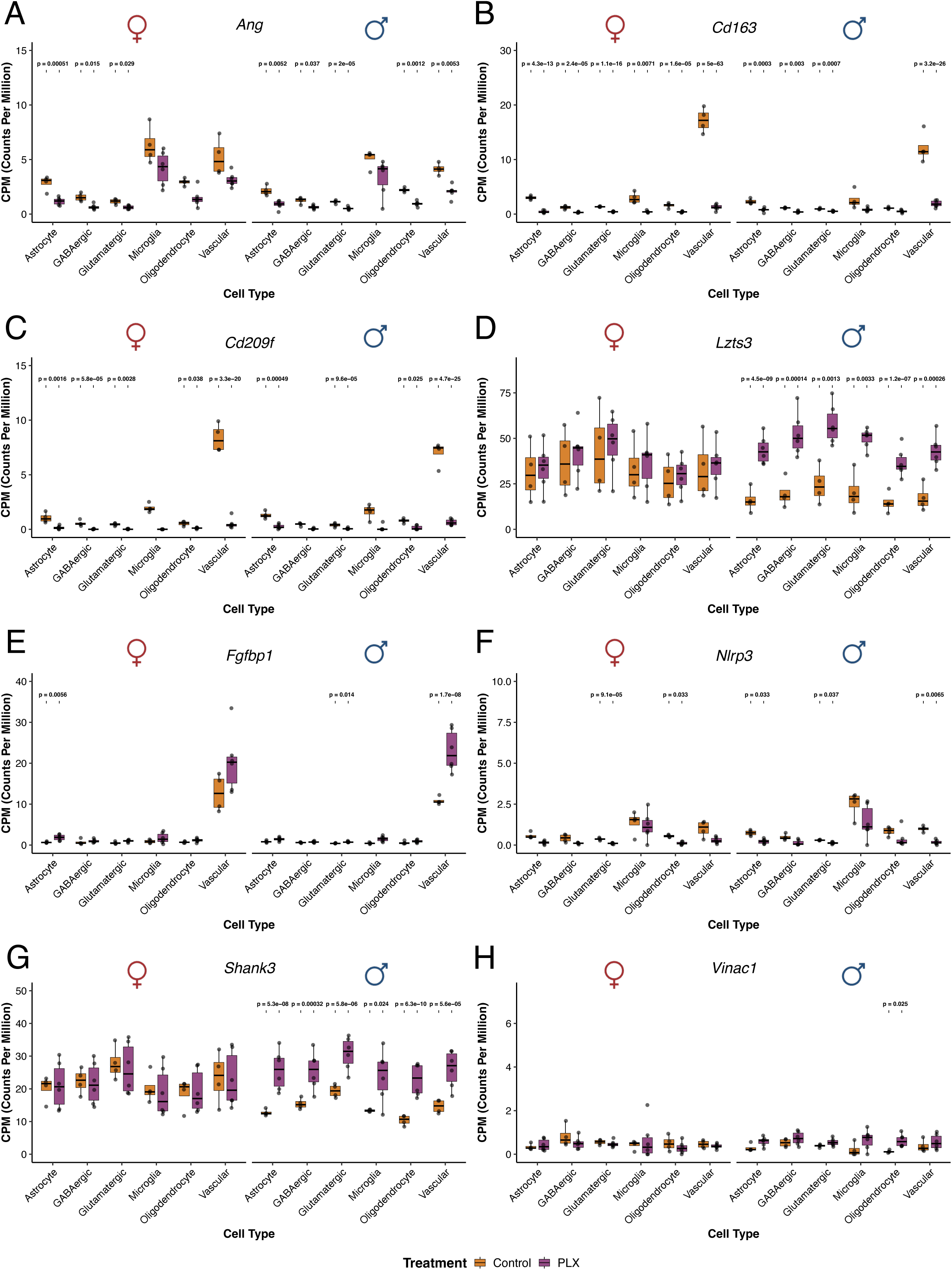
Expression profiles of differentially expressed genes selected by cross-cell-type recurrence or cell-type effect size. Bar plots showing normalized expression (CPM, Counts Per Million) of text-referenced DEGs across six major cell types, separated by sex and treatment. P-values indicate nominal differences between Control and PLX (Welch’s t-test; p-values are displayed on the plot).

**Supplementary figure 7:**
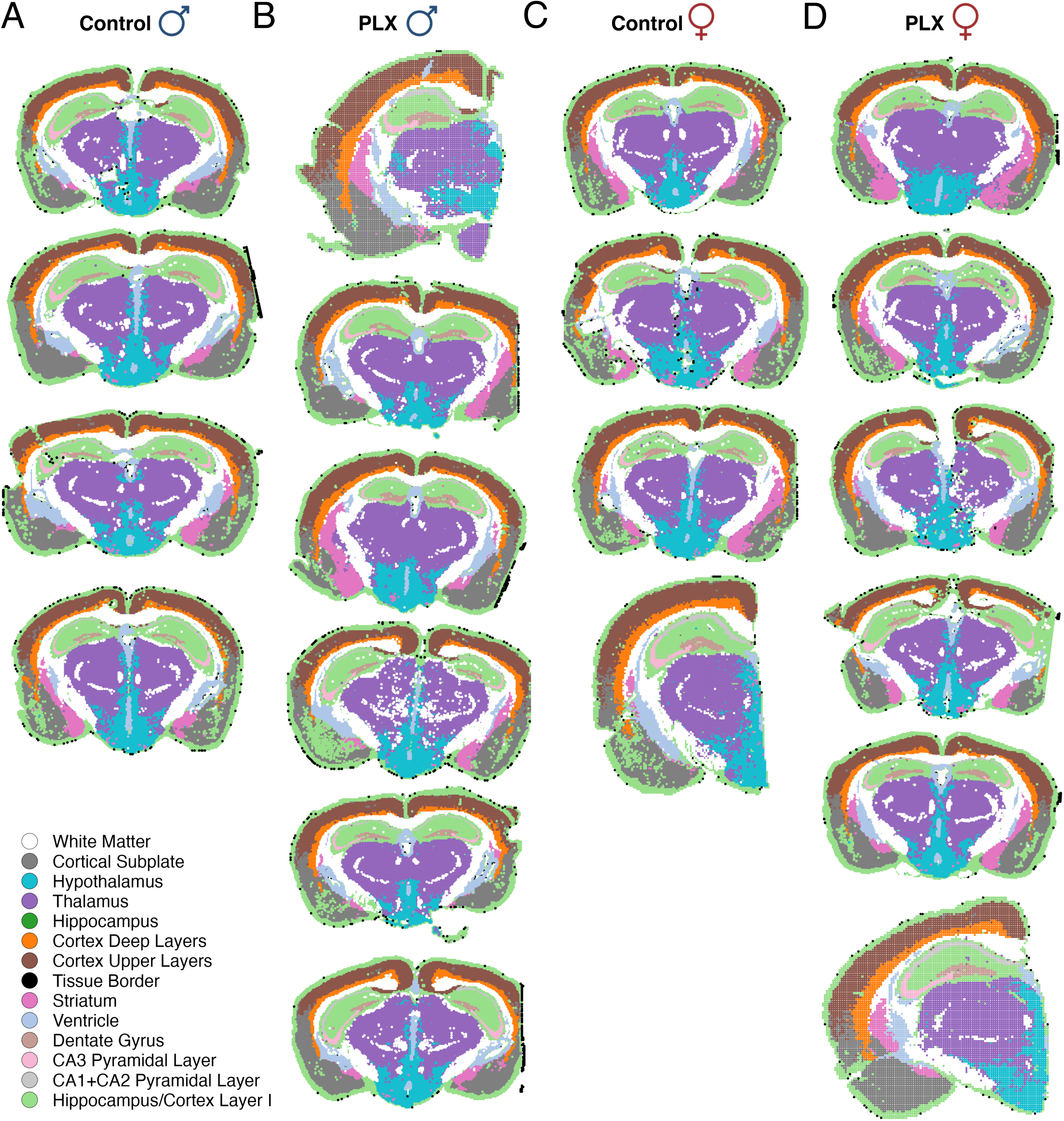
Spatial distribution of anatomical brain regions across all analyzed sections. Spatial maps showing anatomical region annotation across all samples. Each spot is colored by its assigned brain region (legend in the bottom left corner). A. Control male mice (n = 4). B. PLX-treated male mice (n = 6). C. Control female mice (n = 4). D. PLX-treated female mice (n = 6).

**Supplementary figure 8:**
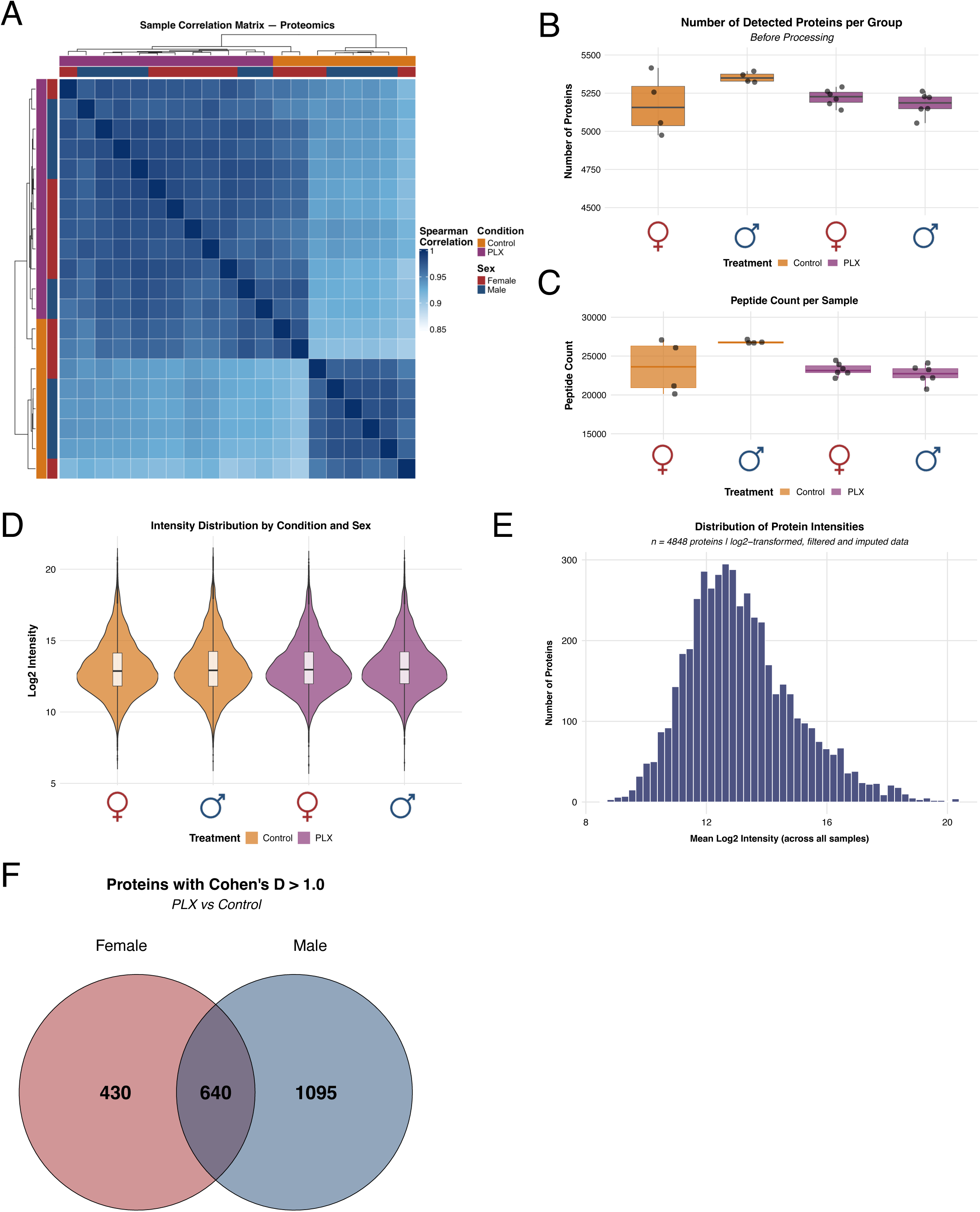
Quality control metrics for bulk proteomics data. A. Heatmap showing Spearman correlation coefficients between all proteomics samples. High correlation values (0.9-1) indicate consistent protein expression profiles across biological replicates. All samples were processed and analyzed in a single batch. B. Box plots showing the number of detected proteins per experimental group before pre-processing. Approximately 5200 proteins were initially detected in each group (before log2-transformation, filtering and imputation), with similar coverage across all conditions. Colors indicate treatment, separated by sex. C. Box plots showing peptide counts per sample across all experimental groups. Median peptide counts range from approximately 20000 – 25000 peptides per sample. Female controls show slightly variable peptide counts, but all groups demonstrate sufficient peptide identification for robust protein quantification. Colors indicate treatment, separated by sex. D. Violin plots showing the distribution of log2-transformed protein intensities across experimental conditions and sex. All groups show similar intensity distributions centered around 12-15 log2 intensity units, with comparable variance. The overlapping distributions confirm protein detection sensitivity across samples and validate the comparability of protein abundance measurements between groups. Colors indicate treatment, separated by sex. E. Histogram showing the mean log2 protein intensities across all samples (n = 4848 proteins after log2-transformation, filtering and imputation). The distribution is normal with a peak around 12 – 14 log2 intensity units. The dataset spans a wide dynamic range of abundances, from low-abundance regulatory proteins to high-abundance structural proteins. F. Venn diagram showing the number of proteins with Cohen’s D > 1.0, with the overlap between sexes. These effect-size-filtered protein sets served as input for the miRNA target-overlap analysis in Fig. 4F.

**Supplementary figure 9:**
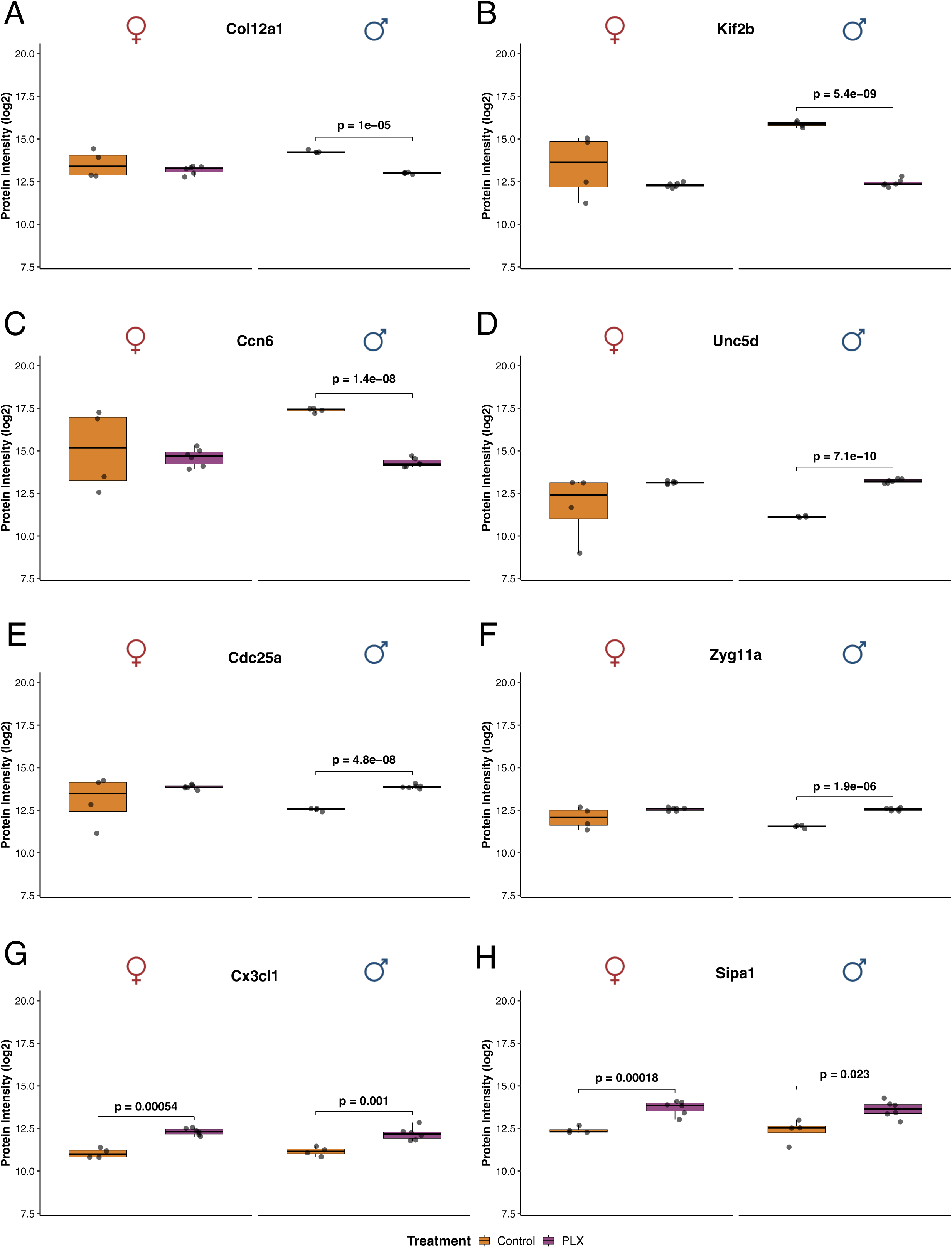
Expression profiles of differentially expressed proteins selected by statistical rank per sex or by membership in the discussed PPI networks. Bar plots showing log2-transformed protein intensities for text-referenced DEPs, separated by sex and treatment. P-values indicate nominal differences between Control and PLX (Student’s t-test; p-values are displayed on the plot).

**Supplementary figure 10:**
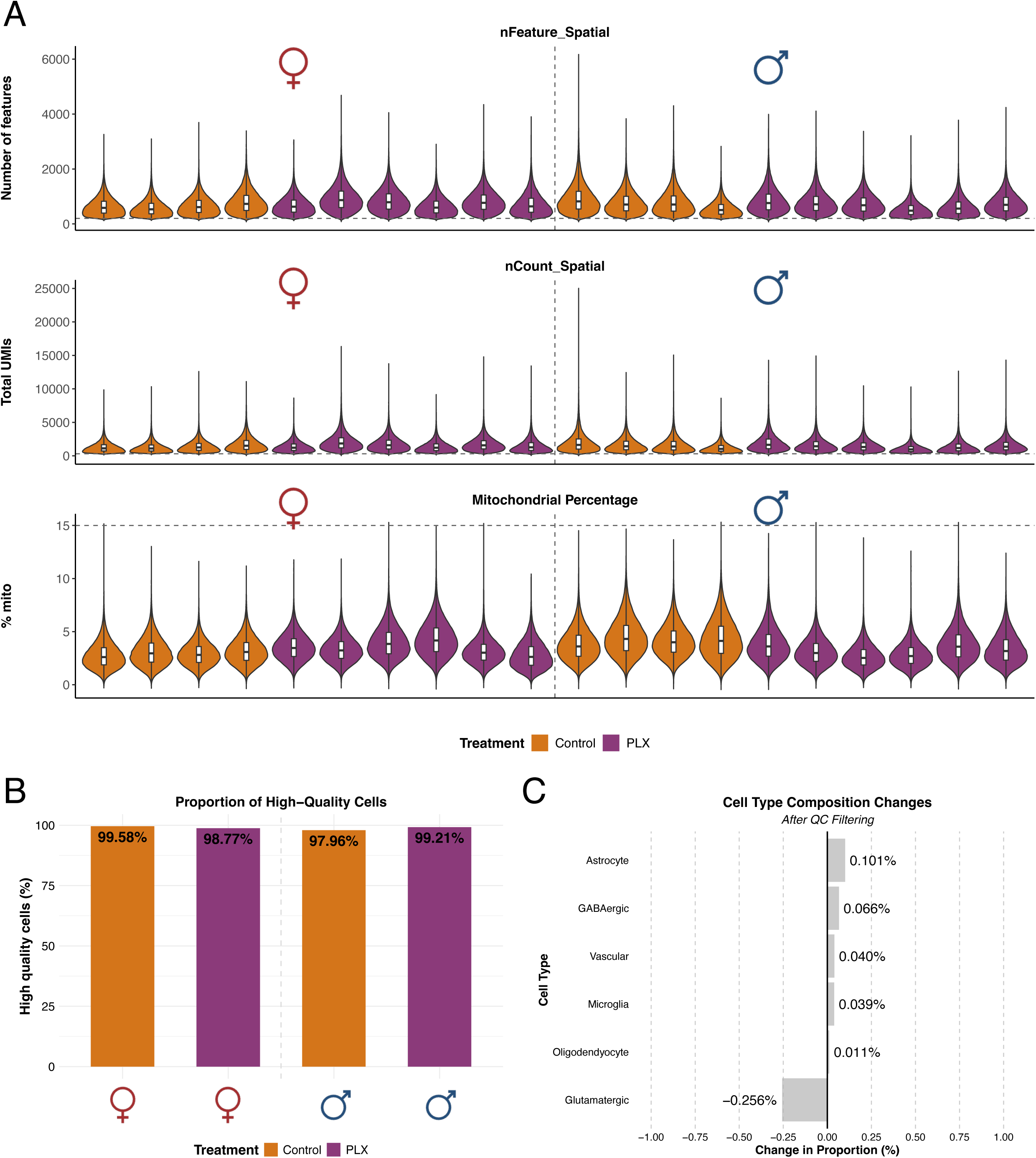
Quality control metrics for the spatial transcriptomics data. A. Violin plots showing quality control metrics for spatial transcriptomics data across all samples. Top panel: number of detected features (nFeature_Spatial) per cell, showing consistent gene detection across conditions. Middle panel: total number of unique molecular identifiers (UMIs; nCount_Spatial) per cell, indicating uniform capture efficiency. Bottom panel: mitochondrial gene percentage (% mito), a metric for cellular stress or RNA quality. Horizontal dashed lines indicate QC filtering thresholds. Colors indicate treatment, separated by sex. B. Bar plot showing the proportion of high-quality cells that are retained after applying quality control filtering thresholds across experimental groups. To assess whether strict QC filtering was necessary, we tested the impact of standard filtering parameters on cell retention. All groups retained >97% of cells, indicating minimal removal of low-quality cells. The uniformly high retention rates across conditions demonstrate that QC filtering removes only minimal low-quality cells. Colors indicate treatment, separated by sex. C. Bar plot showing the cell type composition that occurs after applying stringent QC filtering. Changes in cell type proportions are negligible across all six major cell types. The minimal shifts (<0.3% for all cell types) indicate that QC filtering does not alter the cellular composition of the dataset.

**Supplementary figure 11:**
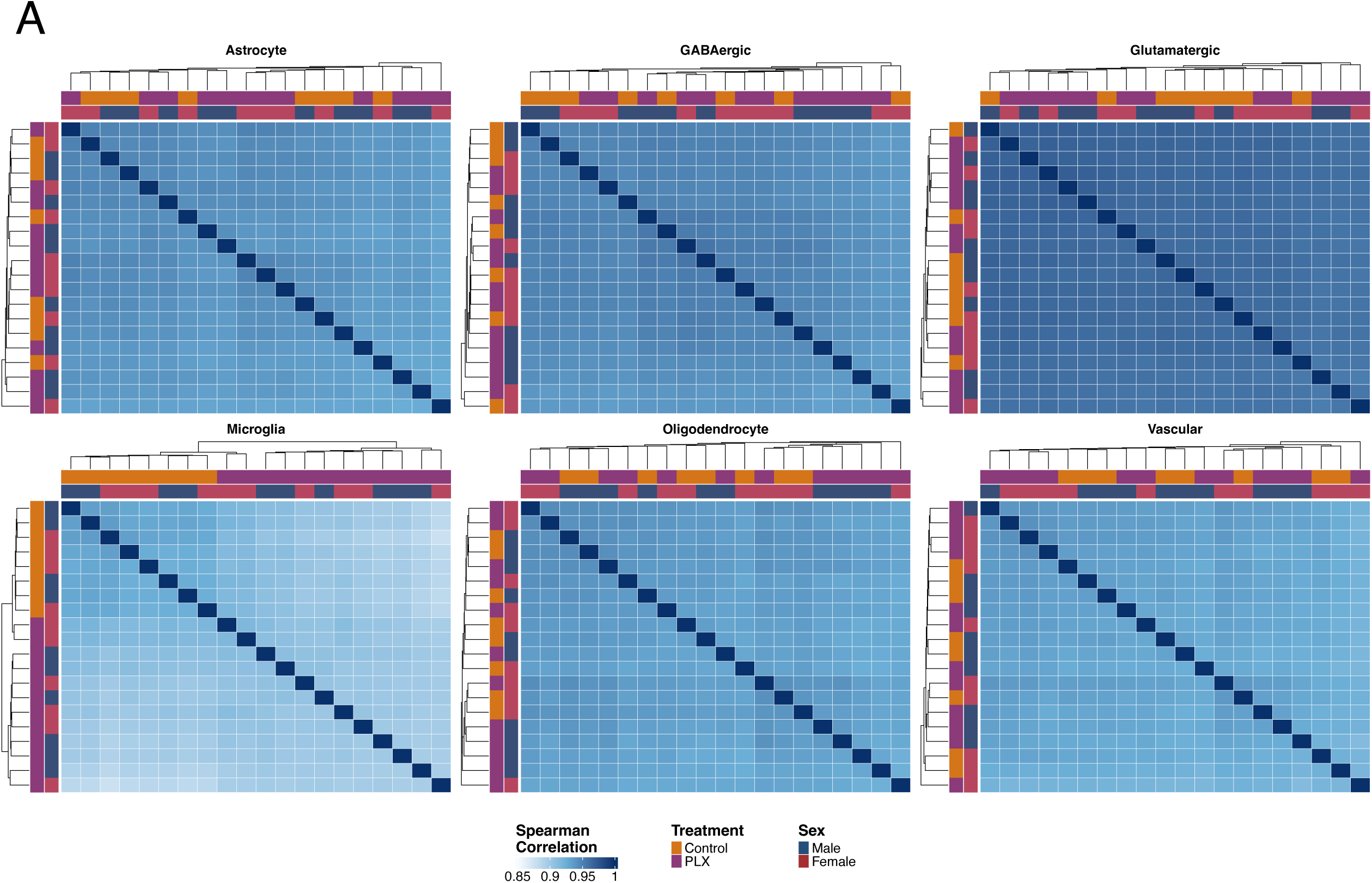
Quality control metrics for the spatial transcriptomics data. A. Heatmap showing Spearman correlation coefficients between samples for each of the six major cell types (astrocytes, GABAergic neurons, glutamatergic, microglia, oligodendrocytes, and vascular cells). High correlation values (0.85 – 1.0) indicate consistent gene expression profiles within cell types across biological replicates. Microglia show slightly lower within-group correlations compared to other cell types. This is likely due to either strong PLX-mediated depletion, resulting in reduced signal, or potential heterogeneity in the remaining microglial subpopulations after CSF1R inhibition.

*Supplementary table 1: Cell type proportion statistics comparing PLX-treated and control mice, stratified by sex*.

Cell type proportions (expressed as percentage of total cells) were calculated from spatial transcriptomics data across all samples. For each of the nine annotated cell types and each sex, the number of samples, mean cell type proportions with standard deviations are reported alongside raw and BH p-values. Microglia were significantly depleted in both sexes (females: -6.1%, adj. p-value = 0.0138; males: -4.1%, adj. p-value = 0.0007), while all other populations remained stable, with the exception of a modest reduction in male ependymal cells.

*Supplementary table 2: Differential expression analysis of detected miRNAs following microglial depletion across brain regions, stratified by sex.*

Complete differential expression results (Control vs. PLX) for all miRNAs, provided as separate sheets for each of the four brain regions and sex. For each miRNA, log2 fold change, fold change, raw p-value, BH adjusted p-value are reported, along with the direction of change and significance calls under both raw (|log2FC| > 1.0, raw p-value < 0.05) and adjusted (|log2FC| > 1.0, adjusted p-value < 0.05) thresholds. The Summary sheet tabulates the number of deregulated miRNAs per regions, sex, and direction under each threshold.

*Supplementary table 3: Cell-type enrichment scores for PLX-deregulated miRNAs from Pomper et al.*^24,30^.

Cell-type enrichment scores (brainstem, all neurons, motor neurons, astrocytes, microglia/immune) are shown for each miRNA deregulated by PLX treatment, alongside the brain regions and sexes affected, and direction of change. ND, no data available. Rows highlighted in pink indicate miRNAs detected in microglial/immune population.

*Supplementary table 4: DEGs per cell type and sex following microglia depletion*.

DEGs identified in six major cell types and both sexes from spatial transcriptomics data (|log2FC| > 1.0, adjusted p-value < 0.05). A Summary sheet reports DEG counts by cell type, sex, and direction; twelve additional sheets (one per cell type x sex) list gene-level statistics. Each gene is categorized into one of three mutually exclusive groups, indicated by both row shading and a corresponding flag column: bona fide DEG retained for downstream analyses (green = bona fide DEGs; red = microglia marker gene (excluded from all cell types); amber = uncharacterized Gm/Rik gene (excluded from all cell types)).

*Supplementary table 5: Cross-cell-type intersections of bona fide DEGs following microglia depletion*.

Bona fide DEGs (as defined in **Supp. Table 4**) were compared across six major cell types to identify shared and cell-type-specific transcriptional responses to PLX treatment, stratified by sex and regulation direction. For each sex/direction combination, genes are grouped by their unique combination of cell types in which they were detected as DEGs, together with the number of cell types and genes per intersection. Intersection set sizes were used as input for UpSet plot visualization (**Fig. 3C**).

*Supplementary table 6: High-effect-size gene/protein pool (Cohen’s D > 1.0) by cell type, and sex*.

Genes and proteins with a large effect size (Cohen’s D > 1.0) between PLX-treated and control samples. Spatial sheets report both sexes together; proteomic sheets are reported separately by sex. Each entry is flagged as a microglia marker gene, an uncharacterized Gm/Rik gene, or a bona fide gene/protein. This high-effect pool defines the input gene and protein for miRNA target enrichment analysis.

*Supplementary table 7: miRNA target genes overlapping with DEGs and DEPs*.

Predicted and validated targets for five candidate miRNAs (miR-142a-3p, miR-142a-5p, miR-146a-5p, miR-223-3p, miR-409-3p); (sourced from TargetScanMouse and miRTarBase). MiRNA targets (defined in **Supp. Table 10**) overlapped with high-effect-size gene/protein (defined in **Supp. Table 6**) are reported here. Overlapping targets were subject to exclusion depending on if it is a microglia marker or an uncharacterized Gm/Rik gene. Red rows = excluded genes.

*Supplementary table 8: DEPs identified by proteomics following microglia depletion*.

DEPs identified in PLX treated versus control brain tissue, separated by sex (|log2FC| > 1.0, raw p-value < 0.05). Proteins are sorted by -log10(p-value) in descending order.

*Supplementary table 9: RNA integrity numbers (RIN) for all samples*.

RIN values measured by Bioanalyzer (Eukaryote Total RNA Nano) for all samples used in small RNA sequencing. All samples yielded RIN > 7.9, with an average RIN value of 8.6.

*Supplementary table 10: Predicted and validated target gene lists for candidate miRNAs*.

Predicted targets (TargetScan Conserved Site Context Scores, weighted context++ score percentile >= 75) and experimentally validated targets (miRTarBase, strong and weak evidence combined) were retrieved for five candidate miRNAs (miR-142a-3p, miR-142a-5p, miR-146a-5p, miR-223-3p, miR-409-3p). No predicted and validated target data were retrieved for miR-433-5p. These target gene lists were used together with the high-effect-size gene/protein pools (defined in **Supp. Table 6**) for miRNA target enrichment analysis.

